# Bidirectional Regulation of Motor Circuits Using Magnetogenetic Gene Therapy

**DOI:** 10.1101/2023.07.13.548699

**Authors:** Santiago R. Unda, Lisa E. Pomeranz, Roberta Marongiu, Xiaofei Yu, Leah Kelly, Gholamreza Hassanzadeh, Henrik Molina, George Vaisey, Putianqi Wang, Jonathan P. Dyke, Edward K. Fung, Logan Grosenick, Rick Zirkel, Aldana M. Antoniazzi, Sofya Norman, Conor M. Liston, Chris Schaffer, Nozomi Nishimura, Sarah A. Stanley, Jeffrey M. Friedman, Michael G. Kaplitt

## Abstract

Here we report a novel suite of magnetogenetic tools, based on a single anti-ferritin nanobody-TRPV1 receptor fusion protein, which regulated neuronal activity when exposed to magnetic fields. AAV-mediated delivery of a floxed nanobody-TRPV1 into the striatum of adenosine 2a receptor-cre driver mice resulted in motor freezing when placed in an MRI or adjacent to a transcranial magnetic stimulation (TMS) device. Functional imaging and fiber photometry both confirmed activation of the target region in response to the magnetic fields. Expression of the same construct in the striatum of wild-type mice along with a second injection of an AAVretro expressing cre into the globus pallidus led to similar circuit specificity and motor responses. Finally, a mutation was generated to gate chloride and inhibit neuronal activity. Expression of this variant in subthalamic nucleus in PitX2-cre parkinsonian mice resulted in reduced local c-fos expression and motor rotational behavior. These data demonstrate that magnetogenetic constructs can bidirectionally regulate activity of specific neuronal circuits non-invasively *in-vivo* using clinically available devices.

**Teaser:** A novel magnetogenetics toolbox to regulate neural circuits *in-vivo*.

## Introduction

Neuromodulation has become a critical tool for both delineating functional brain circuits underlying behaviors in animal studies and treating human patients with circuit disorders (*1*). Electrical stimulation has long been used in pre-clinical studies and deep brain stimulation (DBS) is a standard of care for patients with advanced tremors and Parkinson’s disease (PD) (*2*). Despite the focal effects of electrical stimulation, the need for greater precision in modulating specific circuits or neuronal populations has promoted the development of novel technologies. Optogenetics combines viral delivery of light-sensitive ion channels with regulated light probes for precise millisecond-time scale control of neural activity (*3, 4*), but the microbial origin of the channels and the need for a fiber optic implant can limit applications in animal studies and provide challenges to human translation (*5*). In contrast, chemogenetics delivers genes for modified channels or receptors to allow drug-regulated control of neural activity without an implant (*6*), but the response is dependent on drug diffusion to targeted regions and the time course and longevity of responses rely on the pharmacokinetics of the drug.

Precise spatial and temporal control of neural circuitry without implanted devices could be transformative for both pre-clinical studies and for human therapeutic applications (*7*). *In vivo* magnetic-field-based stimulation to control neural activity has shown promise in mammalian (*5, 7–11*) and non-mammalian systems (*7, 12*). Electromagnetic fields can penetrate tissue and transmit energy to metal/metal oxide particles which in turn gate receptors and ion channels (*13*). Taking advantage of these properties, regulation of several brain structures has been achieved using endogenous or genetically encoded members of the transient receptor potential cation channel subfamily V (TRPV). Tethering of exogenous ferritin or external iron oxide nanoparticles (MNP) (*5*) (*7, 9*) has been shown to lead to alterations of neural activity by magnetic fields *in vitro* and *in vivo*. Previously the use of this system required AAV vectors to introduce fusions of TRPV channels with exogenous subunits of paramagnetic ferritin into the cells or the use of adenovirus vectors to deliver TRPV1 and ferritin cDNA clones which were too large for a single AAV vector.

Here, we report highly cell-specific, temporally precise, remote, and reversible modulation of deep brain structures in awake, freely moving mice using a novel nanobody (Nb) to endogenous ferritin (Ft) fused to the TRPV1 channel (Nb-Ft-TRPV1) encoded by a construct packaged into a single AAV vector. We show that cre-dependent magnetogenetic activation of striatopallidal neurons elicits robust motor freezing in Adora-2a (A2a)-cre mice, similar to that reported using opto- and chemogenetics (*14–16*). The magnetic fields were delivered using standard MRI machines or a commercially available Transcranial Magnetic Stimulation (TMS) device used for human therapy, with a threshold of 180 mT to induce behavior alterations. Neural activation was further confirmed using expression of *c-fos*, positron emission tomography (PET), and *in vivo* fiber photometry to validate magnetogenetic-induced changes of neural activation. Circuit-specific activation of striatopallidal neurons was also achieved in wild type (wt) mice using a retrograde AAV encoding cre recombinase in combination with the cre-conditional magnetogenetic construct. Finally, we achieved significant motor improvements in parkinsonian mice using non-invasive, magnetogenetic inhibition of subthalamic nucleus (STN) neurons with a nanobody-TRPV1 construct modified to block neuronal activity. The approach that is presented provides for a means for inducing bidirectional, real-time regulation of neuronal circuits in freely moving animals in a magnetic field and may have potential for a variety of human therapeutic applications.

## Results

### Generation and validation of ferritin binding nanobodies for magnet stimulation *in vitro*

Previously, we used a bicistronic construct to deliver a TRPV1-anti-GFP nanobody fusion protein together with GFP-tagged ferritin(*9*). However, over-expression of exogenously introduced GFP-tagged mouse ferritin could disrupt cellular iron metabolism and this entire system could not be expressed from a single AAV vector. Therefore, we generated endogenous ferritin-binding nanobodies to bind endogenous ferritin and obviated the need for an exogenous ferritin construct. Our screening assay identified 17 ferritin nanobody (Nb-Ft) clones that bound to both human and mouse ferritin **(Supplementary Fig. 1f & g).** We selected 5 Nb-Ft clones with the highest affinity for human ferritin, as assessed by ELISA **(Fig. 1a)**, for further study. To test their ability to increase intracellular calcium, we generated constructs with each Nb-Ft clone fused to TRPV1 followed by a 2A peptide cleavage and GFP-tagged mouse ferritin. Oscillating magnetic field treatment (465kHz, 30mT) significantly increased calcium-dependent secreted alkaline phosphatase (SEAP) reporter production from HEK-293T cells transfected with Nb-Ft clones 2 or 14 fused to TRPV1 compared to transfected cells without magnet treatment (basal) (**Fig. 1b**, Nb-Ft-TRPV1^Ca2+^ clone 2 -Basal 64.7 ± 13.9 mU/hr, Magnet: 223.0 ± 55.0 mU/hr, p < 0.05, Nb-Ft-TRPV1^Ca2+^ clone 14 - Basal 64.3 ± 19.4 mU/hr, Magnet: 165.5 ± 19.4 mU/hr, p < 0.05).

**Fig. 1.**
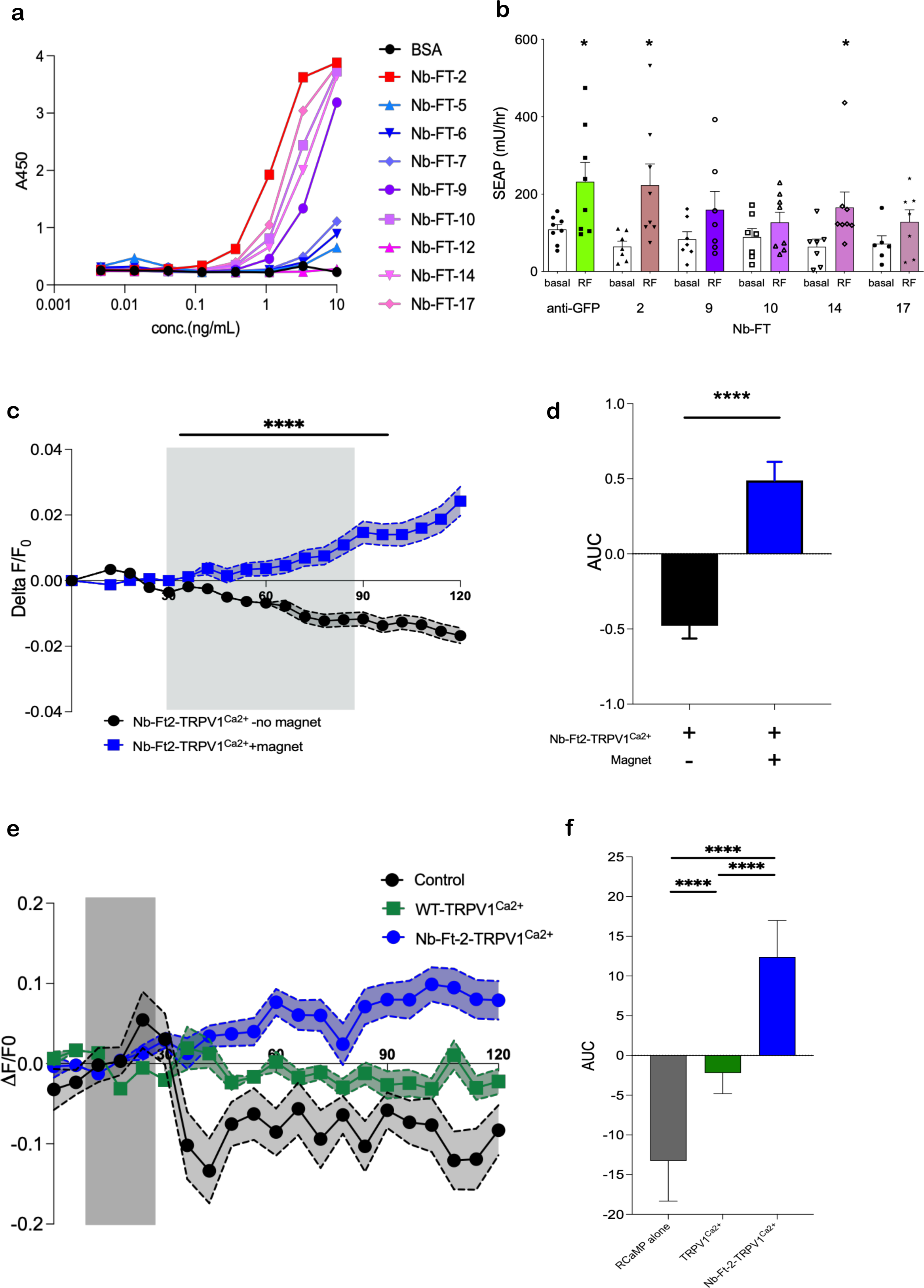
Generation and validation of ferritin binding nanobodies for magnet stimulation. Serial dilutions of nanobodies or BSA (100ul) were incubated on plates coated with human spleen ferritin (1ug/ml) and binding quantified by ELISA after incubation with anti-HA-HRP antibody. **A.** Quantification of anti-ferritin nanobodies from CDR3 groups 6 and 26 binding to human spleen ferritin. **B.** Oscillating magnetic field treatment (465kHz, 30mT) significantly increases calcium-dependent SEAP release from 293T cells transfected with Nb-GFP-TRPV1^Ca2+^, Nb-Ft-2-TRPV1^Ca2+^ and Nb-Ft-14-TRPV1^Ca2+^ and GFP-mFerritin. Data analyzed by two-tailed, unpaired t-test with Welch’s correction. Nb-GFP-TRPV1^Ca2+^, basal vs. RF * p = 0.04, n = 8 & 8; Nb-Ft-2-TRPV1^Ca2+^, basal vs. RF * p = 0.02, n = 7 & 8; Nb-Ft-14-TRPV1^Ca2+^, basal vs. RF * p = 0.04, n = 7 & 8. **C.** Normalized Fluo-4 fluorescence (ΔF/F_0_) in Neuro2A cells expressing Nb-Ft-2-TRPV1^Ca2+^ with (786 cells) or without (1659 cells) magnet treatment. Data were analyzed by two-way ANOVA with Tukey’s multiple comparison test, **** p < 0.0001. **D.** Cumulative change in ΔF/F_0_ of Neuro2A cells expressing Nb-Ft-2-TRPV1^Ca2+^ with (786 cells) or without (1659 cells) magnet treatment. Data were analyzed by Mann Whitney U test **** p < 0.0001. **E.** Changes in RCaMP fluorescence normalized to baseline fluorescence (ΔF/F_0_) with magnet treatment of HEK-293T cells expressing RCaMP alone (54 cells), TRPV1^Ca2+^ (107 cells) or Nb-Ft-2-TRPV1^Ca2+^ (101 cells). Error bars represent mean +/-SEM. **F.** Cumulative change in ΔF/F_0_ with magnet treatment of HEK-293T cells expressing RCaMP alone (54 cells), TRPV1^Ca2+^ (107 cells) or Nb-Ft-2-TRPV1^Ca2+^ (101 cells). AUC was calculated for the period of magnet exposure between 48-180 seconds (after focus adjustment) and analyzed by ordinary one-way ANOVA with Tukey’s multiple comparison test, **** p < 0.0001. Data are shown as mean ± SD.

To evaluate Nb-Ft binding to mouse ferritin, immunoprecipitates (IP) were analyzed from human HEK-293T cells transfected with Nb-Ft-TRPV1 and GFP-tagged mouse ferritin. IPs from cells transfected with any of the 5 clones enriched for human endogenous ferritin. However, only IPs from cells expressing clones 2 and 17 were significant enriched for mouse ferritin compared to expression of GFP-tagged mouse ferritin alone (Mouse ferritin enrichment: Nb-Ft clone 2: 14.5-fold enrichment, clone 9: 1.0 clone 10: 1.1, clone 14: 0.9, clone 17: 6.0 and anti-GFP nanobody: 44.3) (**Supplementary Fig. 2b**). Together, these data revealed that Nb-Ft clone 2 had the highest affinity for human ferritin, greatest enrichment in mouse ferritin after IP, and significantly increased SEAP release with oscillating magnetic field treatment. Thus, anti-ferritin nanobody clone 2 (Nb-Ft-2) was selected for further studies.

We next generated an expression vector with Nb-Ft2 fused to TRPV1^Ca2+^ under the control of a neuronal human synapsin promoter (hSyn-Nb-Ft2-TRPV1^Ca2+^), confirmed cell surface channel expression **(Supplementary Fig. 3a-b)** and function **(Supplementary Fig. 3c-d)**. Magnetic field treatment (190 mT) of murine neuro2A cells expressing Nb-Ft2-TRPV1^Ca2+^ significantly increased in calcium-dependent Fluo-4 fluorescence **(Fig. 1c & d, Supplementary Fig. 4a)** (Peak ΔF/F0: 0.07 ± 0.005 with magnet vs. 0.02 ± 0.001 no magnet, p < 0.0001; AUC: 0.49 ± 0.12 with magnet vs. -0.47 ± 0.08 no magnet, p < 0.0001). Similarly, oscillating magnetic field (465kHz, 30mT) treatment of human HEK-293T cells expressing Nb-Ft-2-TRPV1^Ca2^ significantly increased ΔF/F0 and peak ΔF/F0 RCaMP fluorescence compared to cells expressing wild type TRPV1^Ca2+^ without Nb-Ft-2 fusion or expressing RCaMP alone (control) **(Fig. 1e & f, Supplementary Fig. 4b)** (Peak ΔF/F0: 0.10 ± 0.027 Control, 0.11 ± 0.019 WT-TRPV ^Ca2+^, 0.28 ± 0.034 Nb-Ft-2-TRPV1^Ca2+^, p < 0.0001 Kruskal-Wallis ANOVA. AUC: -13.27 ± 5.05 Control, - 2.19 ± 2.61 WT-TRPV1^Ca2+^, 12.38 ± 4.61 Nb-Ft-2-TRPV1^Ca2+^, *p=0.0195, ****p < 0.0001, Kruskal-Wallis). As a positive control, capsaicin treatment of either wild type WT-TRPV1^Ca2+^or Nb-Ft-2-TRPV1^Ca2+^ in HEK-293T cells significantly increased RCaMP fluorescence **(Supplementary Fig. 3e & f)**. These data indicate that magnetic field treatment increases intracellular calcium in both mouse and human cells expressing the single fusion protein Nb-Ft-2-TRPV1^Ca2+^ (referred to hereafter as Nb-Ft-TRPV1^Ca2+^), encoded by a 2.9 kb cDNA which was used in our AAV vectors for *in vivo* neuronal transduction.

### Magnetogenetic stimulation of striatal indirect pathway alters motor processing *in vivo*

For *in vivo* assessment of functionality of our magnetogenetic construct, we first targeted the indirect pathway using an AAV1/2 vector expressing an HA-tagged, cre-dependent version of the excitatory magnetogenetic Nb-Ft-TRPV1^Ca2+^ cassette, which was flanked by a double floxed inverted open reading frame (DIO) under the control of the JET promoter (AAV1/2-JET-DIO-Nb-Ft-TRPV1^Ca2+^-HA). The cre-dependent Nb-Ft-TRPV1^Ca2+^ AAV was injected bilaterally into the dorsal striatum of transgenic mice expressing cre-recombinase in indirect spiny projection neurons (iSPN) expressing dopamine type 2 (D2) receptors (A2a-cre mice). Immunostaining for HA or RFP (for the AAV1/2-JET-DIO-mCherry control group) demonstrated efficient gene expression in the dorsal striatum **(Fig. 2a)**. Histological analysis of transcript expression using RNAScope confirmed that AAV transcripts co-localized with D2-type neurons and had minimal co-localization with D1-type neurons (D1=3.9% vs. D2=100%, p<0.001) **(Fig. 2b & c)**; the small percentage of D1 transduction is likely due to double positive SPN for D1 and D2 RNA probes **(Supplementary Fig. 5a)**. To determine if this new magnetogenetic construct could functionally activate iSPNs *in vivo*, we first examined motor activity after placing animals in a direct magnetic field (DMF) **(Fig 2d)**. Parkinson’s disease is associated with increased iSPN activity and optogenetic activation of these neurons in normal mice results in rapid freezing (*14*). Bilateral DMF stimulation (500 mT-1.3 T) using a 3T MRI magnet significantly decreased activity compared to baseline, with activity returning to normal after removal from the DMF **(Fig. 2e)**. There was also a marked increase in the time spent freezing (n=10, Nb-Ft-TRPV1^Ca2+^ + DMF=20.3 ± 2.8 vs mCherry +DMF=1.2 ± 0.69 sec, p<0.0001), and a decrease of ambulation (n=10, Nb-Ft-TRPV1^Ca2+^ + DMF= 23.0 ± 5.1 vs mCherry +DMF= 44.7 ± 1.9 sec, p=0.001) during DMF application in the Nb-Ft-TRPV1^Ca2+^ group placed in a magnetic field compared to the mCherry group **(Fig. 2f & g)**. Latency time from DMF exposure until onset of gait freezing was commonly observed between 40 to 60 seconds following exposure to the magnetic field, whereas mice returned to motor baseline immediately post-DMF (**Supplementary video 1**). The mCherry group did not show any motor change upon DMF exposure (**Supplementary video 2**). Minor differences in transduction efficiency had no correlation with freezing of gait **(Supplementary Fig. 5b)**.

**Fig. 2.**
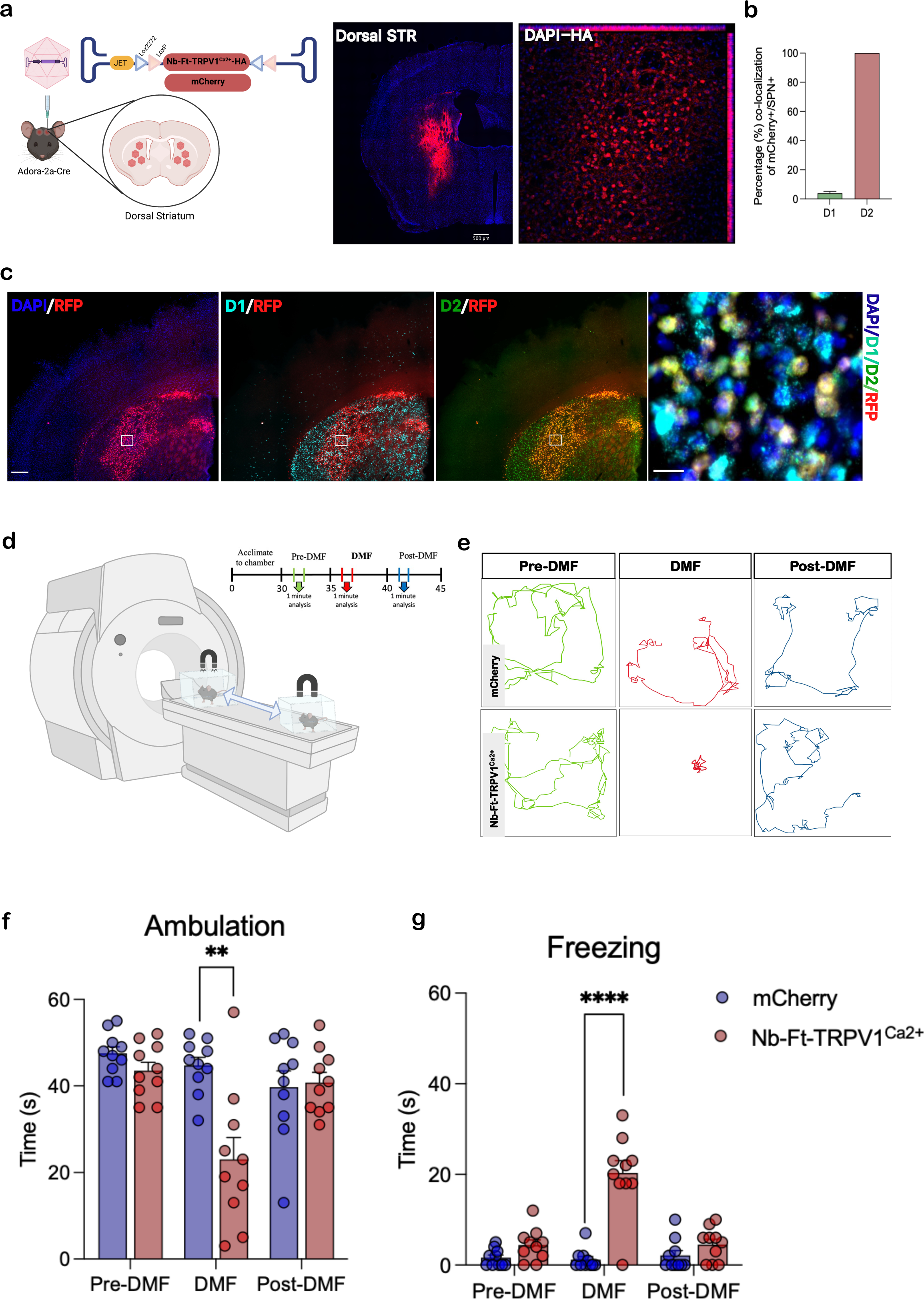
Selective viral-mediated expression of Nb-Ft-TRVP1^Ca2+^ in striatal iSPNs elicit parkinsonian motor behavior. **A.** Schema of the double-floxed Cre-dependent AAV vector expressing the Nb-Ft-TRVP1^Ca2+^ or mCherry under the control of the JET promoter, and immunostaining for HA in Adora-2A (A2a)-Cre mice demonstrates selective expression of the AAV-Nb-Ft-TRVP1^Ca2+^ in dorsal striatum. **B.** Selective transduction of D2-type iSPN with RNA in-situ hybridization (ISH). Co-localization of D2-type (green) iSPN with mCherry (red). **C.** Protocol and Schema for direct magnetic field (DMF) neural activation to assess motor behavior. **D.** Example of altered motor activity during bilateral striatal pre-DMF (green), DMF (red) and post-DMF (blue) stimulation; individual lines represent the path of a single mouse. Effect of DMF stimulation on **E.** Ambulation bout duration, and **F.** Freezing of gait in mCherry (blue bars/dots, n=10) and Nb-Ft-TRVP1^Ca2+^ (red bars/dots, n=10) mice. Error bars show SEM. ** represents p value < 0.01, **** represents p value <0.0001 with two-tailed, unpaired t-test with Welch’s correction .

To determine the threshold for magnetic field strength required to regulate striatal cellular activity, we compared two different field gradients, 500 mT-1.3 T (High DMF titration) **(Supplementary Fig. 6a)** and 100 mT-270 mT (Low DMF titration) as determined by placing a gaussmeter in different locations in the cage at different distances from the 3T magnet **(Fig. 3a)**. After mapping the magnetic field strength in the behavioral boxes **(Fig. 3b & Supplementary Fig. 6b)**, we then recorded the locomotor activity in animals expressing the magnetogenetic construct. The activity heatmap in the presence of the Low DMF revealed a consistent decrease of activity only when the mice receiving Nb-Ft-TRPV1^Ca2+^ were in proximal sections of the behavior box with a field strength at or above 180 mT **(Fig. 3b)**. This contrasted with animals exposed to the High DMF which showed reduced activity across the entire box **(Supplementary Fig. 6c)**. Thus, while the freezing time was consistent throughout the box in the mice in the High DMF titration **(Supplementary Fig. 6d)**, in the Low DMF titration there was a significant increase in freezing time (n=5, p<0.05) only in the regions with >180 mT field strength **(Fig. 3b)**. This threshold is consistent with a prior (*10*) report which used an earlier generation magnetogenetic system consisting of separate TRPV1 and ferritin cDNAs to demonstrate that a DMF source of ∼0.2 T was sufficient to regulate deep brain circuits involved in food intake and glucose homeostasis. To further evaluate whether a magnetic field gradient over 180 mT is sufficient to induce a robust motor phenotype, we repeated our behavioral assessment at low DMF and high DMF **(Fig. 3c)**. Consistent with the prior results, we found that mice expressing Nb-Ft-TRPV1^Ca2+^ stayed at a relatively fixed location during both Low and High DMF application while control mice expressing mCherry continued to move normally during DMF conditions **(Fig. 3d & e)**. Indirect pathway activation with both High and Low DMF application decreased the time spent ambulating (n=5-8, Low DMF Nb-Ft-TRPV1^Ca2+^= 15.75 ± 2.4 vs mCherry= 36 ± 3.48, p<0.001; No DMF Nb-Ft-TRPV1^Ca2+^= 35.63 ± 2.16 vs mCherry= 46.6 ± 2.54, p<0.01; High DMF Nb-Ft-TRPV1^Ca2+^= 10 ± 2.89 vs mCherry= 44.2 ± 2.31, p<0.001), and increased time spent freezing (n=5-8, Low DMF Nb-Ft-TRPV1^Ca2+^= 20.38 ± 2.4 vs mCherry= 5.4 ± 0.74, p<0.01; No DMF Nb-Ft-TRPV1^Ca2+^= 8.13 ± 1.92 vs mCherry= 3.2 ± 1.46, p>0.05; High DMF Nb-Ft-TRPV1^Ca2+^= 25.5 ± 4.01 vs mCherry= 4 ± 0.83 p<0.01). **(Fig. 3f & g)**.

**Fig. 3.**
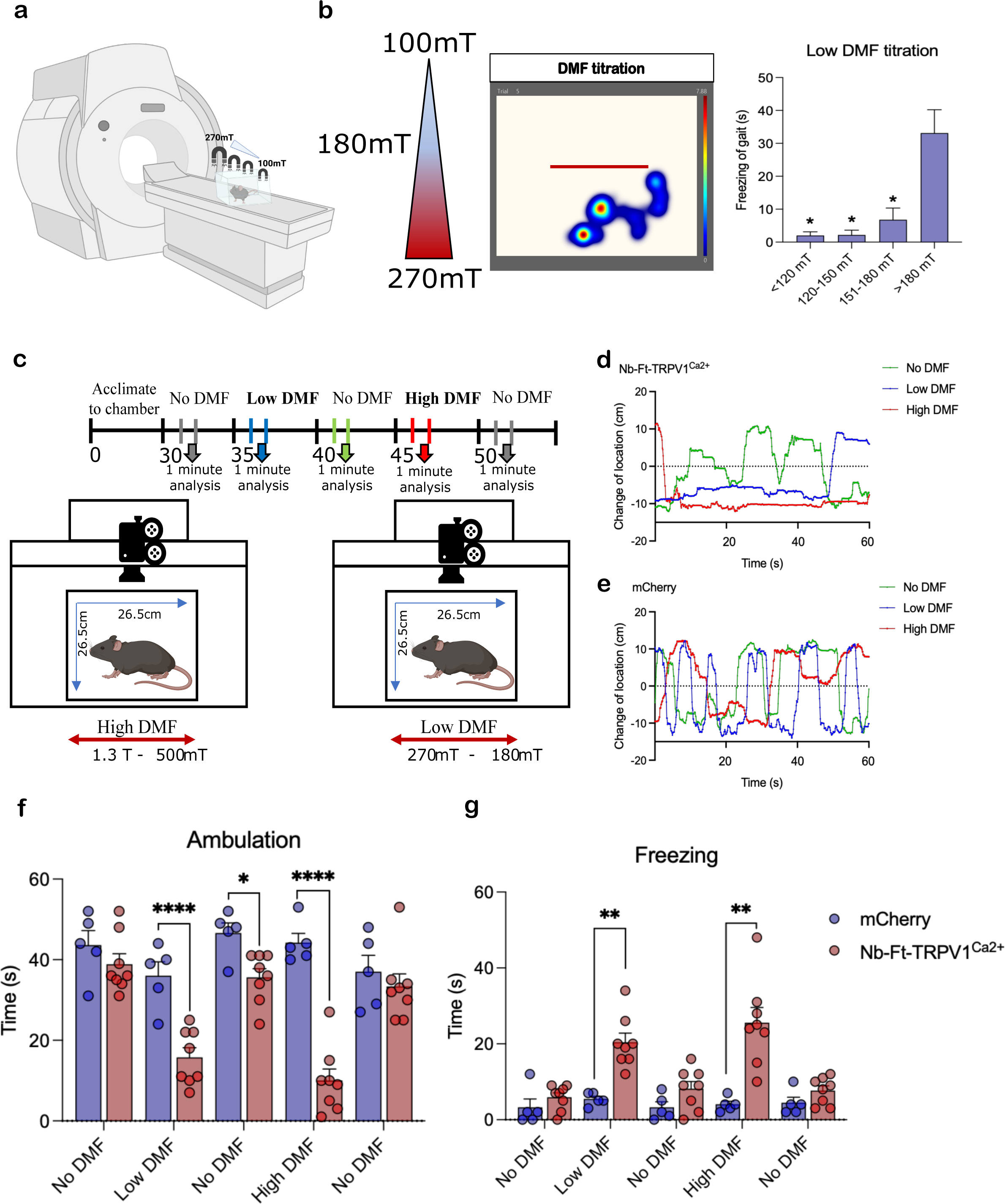
Nb-Ft-TRPV1^Ca2+^ in D2 iSPNs stimulated with high DMF and low DMF treatment produce freezing of gait. **A.** Schema for low direct magnetic field (DMF) titration. **B.** Magnetic field strength and individual activity heatmap example of motor activity during bilateral striatal Low DMF titration in Nb-Ft-TRPV1^Ca2+^ expressing mice. Freezing of gait in different magnetic field gradients in the Low DMF titration ranges (blue bars, n=5) in Nb-Ft-TRPV1^Ca2+^ expressing mice. **C.** Protocol and schema for High DMF and Low DMF stimulation in A2a-Cre mice injected with the double-floxed Cre-dependent AAV vector expressing the Nb-Ft-TRPV1^Ca2+^ or mCherry under the control of the JET promoter. Representative tracking data of change of location in **D.** Nb-Ft-TRPV1^Ca2+^ and **E.** mCherry A2a-Cre mice during High DMF (red line), Low DMF (blue line), and No DMF (green line) treatment. Effect of High DMF and Low DMF stimulation on **F.** Ambulation and **G.** Freezing of gait in Nb-Ft-TRPV1^Ca2+^ (red dot/bars, n= 8) and mCherry (blue dots/bars, n=5). Error bars show SEM. * represents p value < 0.05, ** represents p value < 0.01, **** represents p value < 0.0001 with two-tailed, unpaired t-test with Welch’s correction .

### Magnetogenetic stimulation increases striatal neuronal activity *in vivo*

We next assessed whether the changes in locomotor activity were correlated with changes in neural activity. Previously the immediate early gene *c-fos* was shown to be increased in *vitro* and *in vivo* in cells expressing TRPV1 and ferritin in a magnetic field (*8, 10, 11*). Consistent with this, a significantly higher proportion of striatal neurons expressing Nb-Ft-TRPV1^Ca2+^ showed c-*fos+* expression compared to mCherry controls following exposure exposed to the MRI magnet (500 mT-1.3 T) (n=5, Nb-Ft-TRPV1^Ca2+^=23.65 ± 3.49 vs mCherry= 5.79 ± 0.77 %, p<0.01) **(Fig. 4a & b)**. There was a correlation between *c-fos* expression and freezing (n=4-5, Pearson’s r=0.39, two-tailed n.s.) with the Nb-Ft-TRPV1^Ca2+^ and mCherry mice clustering at opposite extremes of the correlation matrix **(Fig. 4c)**. The data are consistent with there being a robust activation of the indirect pathway neurons, with relatively low variability between the expression of gene activity-induced makers, and the freezing phenotype.

**Fig. 4.**
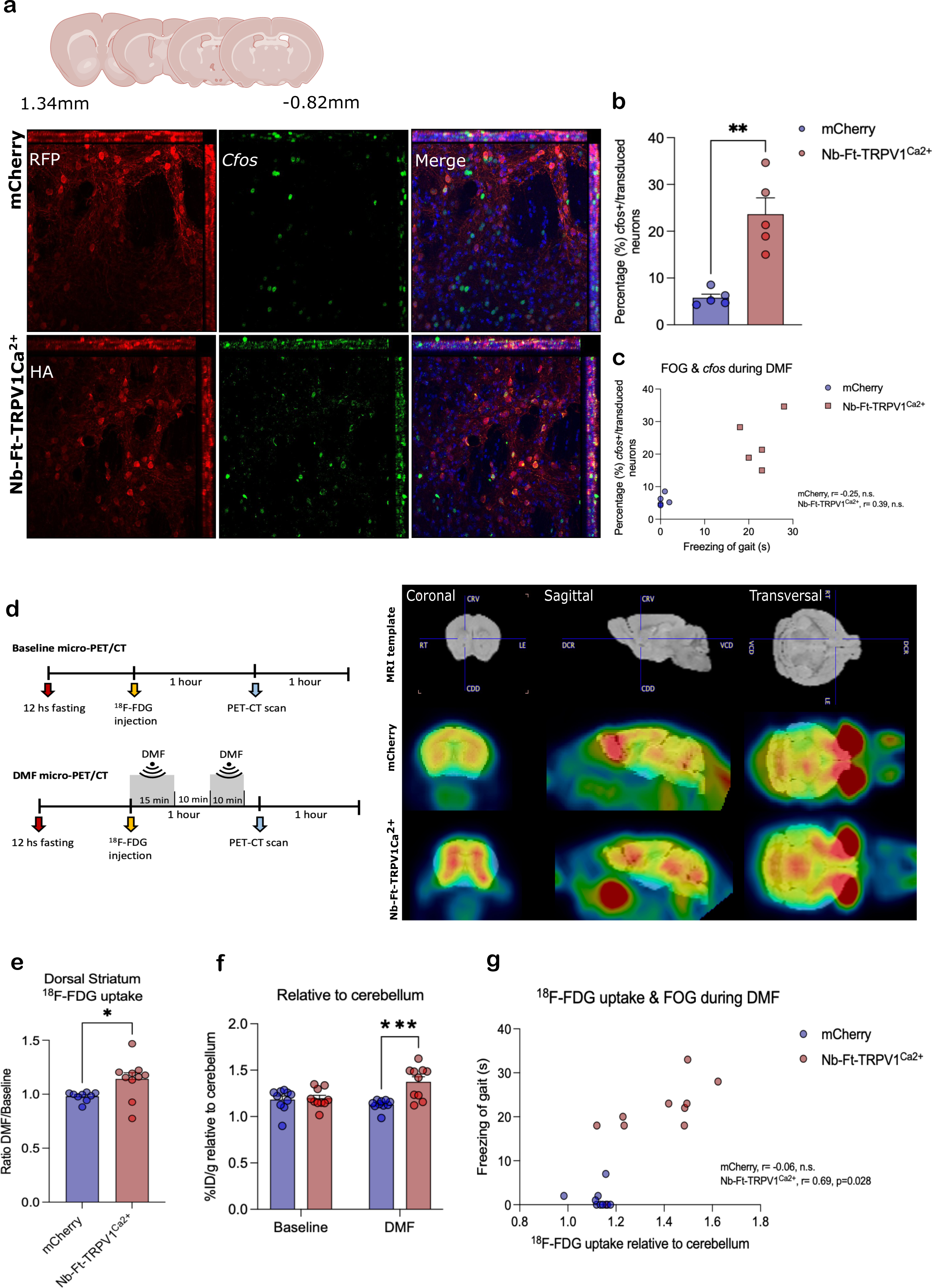
Nb-Ft-TRPV1^Ca2+^ increases *c-fos* expression and cerebral metabolism upon DMF treatment. **A.** Immunostaining for *c-fos* (cyan) and HA or RFP (red) of dorsal striatum of A2a-Cre mice euthanized 1 hour following 15 minutes of DMF exposure. **B.** Percentage (%) of *c-fos* positive (+) neurons over total number of iSPNs expressing HA or RFP (n=5 mice per group/3 sections per mice). **C.** Correlation between *c-fos* expression and freezing of gait during DMF exposure (n=5 mice per group). **D.** Protocol and schema for 18-FDG micro-PET/CT scans procedure and analysis with representative micro-PET/CT imaging of A2a-Cre mouse brains expressing the Nb-Ft-TRPV1^Ca2+^ or the mCherry control virus following DMF exposure. 18-FDG uptake in dorsal striatum calculated as the percentage of injected dose/weight (%ID/g), **E.** DMF over baseline ratio. **F.** DMF over baseline normalized to cerebellum, in mCherry (blue bars/dots, n=8) and Nb-Ft-TRPV1^Ca2+^ (red bars/dots, n=8) mice. **G.** Correlation between DMF over baseline 18-FDG uptake ratio and freezing of gait during DMF exposure (n=10). Error bars show SEM. * represents p value < 0.05, ** represents p value < 0.01, *** represents p value < 0.001 with two-tailed, unpaired t-test with Welch’s correction. Anti-HA staining to detect Nb-Ft-TRVP1^Ca2+^ expression, anti-RFP staining to detect mCherry expression.

To further assess striatal activity after magnetogenetic activation living animals, we used positron emission tomography (PET) with [^18^F] fluorodeoxyglucose (^18^F-FDG), which measures glucose utilization as a reflection of underlying neural activity (*18*). Micro-PET-computed tomography (CT) was performed in A2a mice expressing Nb-Ft-TRPV1^Ca2+^ fasted for 12 hrs. vs mice receiving an mCherry in the striatum construct. ^18^F-FDG was administered to the two groups first without DMF exposure, followed 3 days later by PET imaging following the application of DMF **(Fig. 4d left panel)**. DMF exposure significantly increased striatal uptake of ^18^F-FDG, expressed as the ratio of uptake at baseline prior to DMF exposure to the magnetic field to the signal after placement in a magnetic field. The ratio of the PET signal was significantly greater in Nb-Ft-TRPV1^Ca2+^ group compared to mCherry controls (n=9-10, Nb-Ft-TRPV1^Ca2+^= 1.14 ± 0.06 vs mCherry= 0.98 ± 0.001, p<0.05) **(Fig. 4d right panel & e)**. To account for individual variability, we also quantified the percentage of the ^18^F-FDG dose that was injected dose gram (%ID/g) in the striatum compared to uptake in cerebellum, with and without DMF treatment. DMF exposure significantly increased ^18^F-FDG uptake in the striatum in mice with Nb-Ft-TRPV1^Ca2+^ expression in D2 neurons relative to mCherry controls (n=10, Nb-Ft-TRPV1^Ca2+^= 1.37 ± 0.06 vs mCherry= 1.19 ± 0.03 %ID/g, p<0.001) **(Fig. 4f)**. No change in the PET signal was seen in cerebellum. Since clinically ^18^F-FDG PET-CT is used to monitor Parkinson’s disease progression and can be correlated with motor features (*18*), we reasoned that the levels of striatal glucose uptake might correlate with the time mice spent freezing during DMF application. Consistent with this, the data showed a significant positive correlation (n=7-9, Pearson’s r=0.69, two-tailed p<0.05) between striatal glucose uptake and freezing time in the Nb-Ft-TRPV1^Ca2+^ expressing mice **(Fig. 4g)**, with the most marked increase in the time spent freezing observed in animals with a ^18^F-FDG uptake over 1.4 using the cerebellum as a reference.

Finally, we used genetically encoded calcium indicators to measure magnetogenetic-induced neural activity (*5, 7, 11, 12*). To selectively record D2 iSPN calcium transients during pre-DMF, DMF, and post-DMF periods, we bilaterally injected a cre-dependent excitatory magnetogenetic vector (AAV1/2-JET-DIO-Nb-Ft-TRPV1^Ca2+^ -HA) together with a cre-dependent calcium indicator (AAV9-Syn-DIO-GCaMP6s) and implanted an optical fiber in the dorsal striatum of A2a-cre mice **(Fig. 5a & b)**. This required the use of optical fibers that did not contain any metal which would heat in the presence of a magnetic field (**Fig. 5c**). Nb-Ft-TRPV1^Ca2+^ expressing mice again showed motor freezing in response to DMF that was reflected with an increased ΔF/F striatal GCaMP fluorescence signal (**Fig. 5f)**, which was not seen in mCherry expressing mice **(Fig. 5d)**. Mean GCaMP6 fluorescence (ΔF/F) percentage aligned to Pre-DMF, DMF, and Post-DMF revealed a marked increase of fluorescence during DMF stimulation in the Nb-Ft-TRPV1^Ca2+^ group **(Fig. 5g)** but not in the mCherry group **(Fig. 5e)**. Calcium transients in D2 iSPNs demonstrated by an increase in GCaMP6 fluorescence (ΔF/F) were significantly increased by DMF treatment in the Nb-Ft-TRPV1^Ca2+^ but not in the mCherry expressing mice (n=3-4, Nb-Ft-TRPV1^Ca2+^ + DMF= 27.91 ± 8.24 vs mCherry +DMF= -1.37 ± 1.05%, p<0.05) **(Fig. 5i)**. In parallel, DMF treatment significantly increased time spent freezing in Nb-Ft-TRPV1^Ca2+^ compared to mCherry controls (n=3-4, Nb-Ft-TRPV1^Ca2+^ + DMF= 29.33 ± 7.3 vs mCherry +DMF= 3.5 ± 1.3, p<0.05) **(Fig. 5h)** confirming the functional efficacy of magnetogenetic activation. Consistent with the *c-fos* expression level and the PET studies, there was a positive correlation between the change in fluorescent signal and the extent of the motor inhibition (n=3-4, Pearson’s r=0.60, two-tailed n.s.), with the experimental and control groups mostly clustering at opposite ends of the correlation matrix **(Fig. 5j)**. This combined data demonstrates that the fused ferritin nanobody-TRPV1 channel, expressed from a single AAV in striatal D2 neurons, resulted in activation of transduced iSPNs which then caused the resulting motor freezing.

**Fig. 5.**
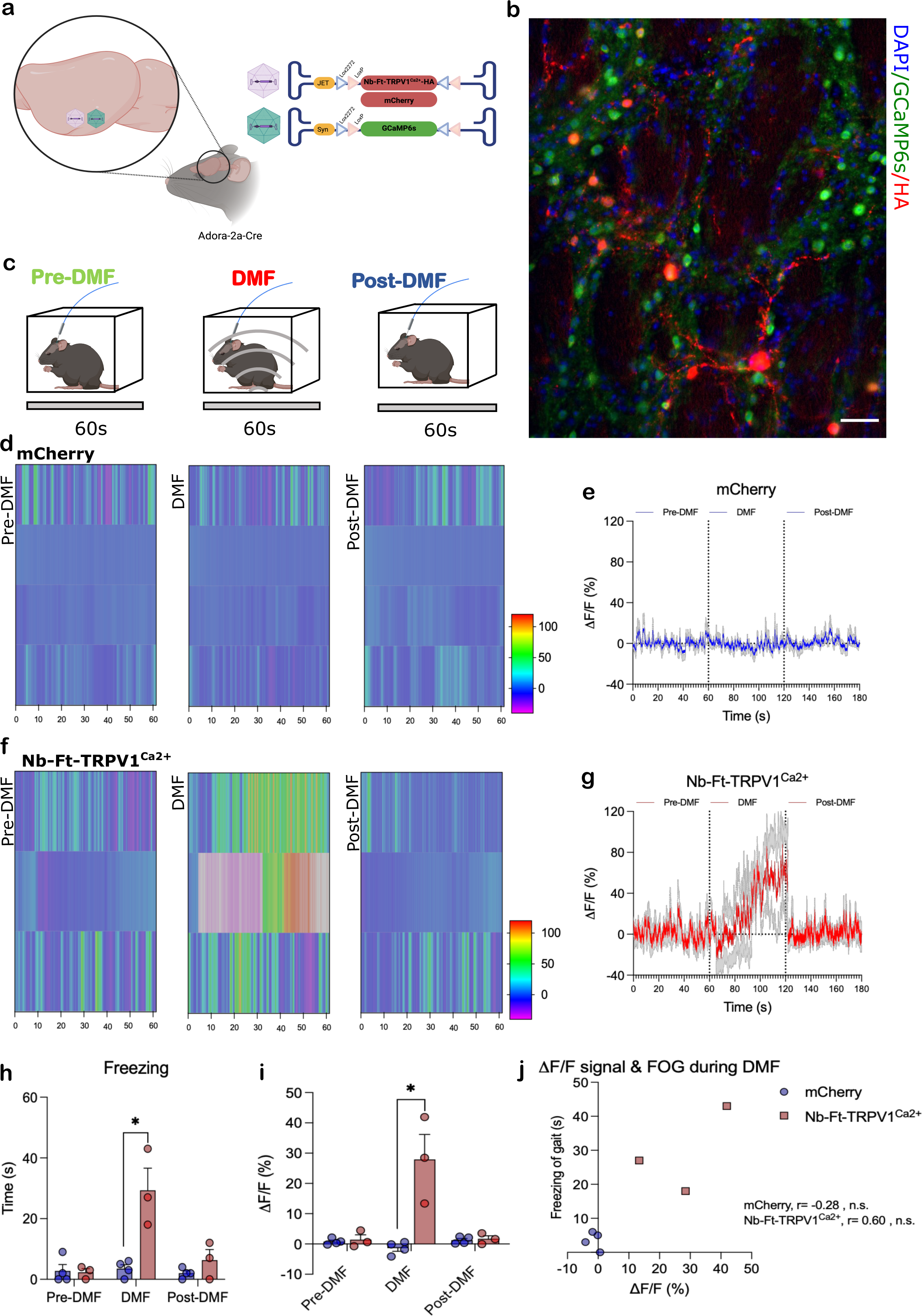
Nb-Ft-TRPV1^Ca2+^ increases calcium transients in dorsal striatum. **A.** Schema of the double-floxed Cre-dependent AAV vector expressing the Nb-Ft-TRPV1^Ca2+^ or mCherry under the control of the JET promoter (red) combined with double floxed Cre-dependent AAV vector expressing GCaMP6s under the control of the Syn promoter (green). Behavioral methodology to assess calcium transients during pre-DMF (green), DMF (red), and post-DMF (blue) treatment. **B.** Activity trace example of altered motor activity during bilateral striatal pre-DMF (green), DMF (red), and post-DMF (blue) stimulation in an Nb-Ft-TRPV1^Ca2+^ mouse with heatmap of GCaMP fluorescence aligned to pre-DMF (left), DMF (middle), and post-DMF (right) of the same animal shown in activity trace example expressing the Nb-Ft-TRPV1^Ca2+^ vector. **C.** Mean fluorescence aligned to pre-DMF, DMF, and post-DMF in Nb-Ft-TRPV1^Ca2+^ group (n=3). **D.** Activity trace example of altered motor activity during bilateral striatal pre-DMF (green), DMF (red), and post-DMF (blue) stimulation in an mCherry mouse with heatmap of GCaMP fluorescence aligned to pre-DMF (left), DMF (middle), and post-DMF (right) for the same animal shown in activity trace example expressing the mCherry vector. **E.** Mean fluorescence aligned to pre-DMF, DMF, and post-DMF in the mCherry group (n=4) **F.** Freezing of gait (FOG), **G.** Average delta (Δ) F/F (%) during pre-DMF, DMF, and post-DMF stimulation. **H.** Correlation between ΔF/F (%) calcium transients and FOG during DMF treatment. Error bars show SEM. * represents p value < 0.05 with two-tailed, unpaired t-test with Welch’s correction.

### Transcranial magnetic stimulation (TMS) increases Nb-Ft-TRPV1^Ca2+^ mediated calcium transients and elicits parkinsonian phenotype

While the MRI magnet was highly effective for demonstrating and characterizing the magnetogenetic effect of the new construct on neurons within the brain, the feasibility of translating into human application a technology which requires proximity to an MRI for therapeutic efficacy is limited. To evaluate the potential of a more clinically relevant technology, we repeated the studies above using a commercial transcranial magnetic stimulation (TMS) system which is FDA approved for depression and other applications (*19*) **(Fig. 6a)**. Similar to the effect of an MRI machine, TMS application at 20% output capacity using 50 twin pulses per second with 1 second inter-stimulus intervals decreased motor ambulation in Nb-Ft-TRPV1^Ca2+^ expressing mice compared with mCherry controls **(Fig. 6b)**. Analysis of motor parameters including distance moved **(Fig. 6c & d)**, activity **(Supplementary Fig. 7a & b)** and change of location **(Supplementary Fig. 7d & e)** was consistent with activation of D2 iSPN in Nb-Ft-TRPV1^Ca2+^ expressing mice compared to mCherry controls. Average distance moved and activity percentage during TMS application was significantly decreased in the Nb-Ft-TRPV1^Ca2+^ expressing mice from baseline (n=5, Nb-Ft-TRPV1^Ca2+^ + Baseline= 32.75 ± 4.15 vs Nb-Ft-TRPV1^Ca2+^ + TMS= 19.98 ± 1.57, p<0.05) and compared to mCherry controls (n=4-5, Nb-Ft-TRPV1^Ca2+^ + TMS= 19.98 ± 1.57 vs mCherry + TMS= 30.94 ± 2.71, p<0.01) **(Fig. 6e, Supplementary Fig. 7c)**. Again, using targeted GCAMP expression and fiber photometry, TMS treatment produced rapid generation of fluorescent positive spikes in the striatum of Nb-Ft-TRPV1^Ca2+^ expressing mice while TMS treatment showed a sustained decrease in GCaMP6 fluorescence in mCherry-expressing controls **(Fig. 6f & g)** (*20*). Fluorescence (ΔF/F) quantification showed a significant increase in the Nb-Ft-TRPV1^Ca2+^ expressing mice compared to their own baseline (n=3, Nb-Ft-TRPV1^Ca2+^ + TMS=14.41 ± 4.15 vs Nb-Ft-TRPV1^Ca2+^ + Baseline= -2.38 ± 0.81%, p<0.05) and compared with mCherry mice during TMS application (n=3-4, Nb-Ft-TRPV1^Ca2+^ + TMS= 14.41 ± 4.15 vs mCherry + TMS= -15.81 ± 5.61%, p<0.01) **(Fig. 6h)**. These studies confirm that, as was seen with MRI generated magnetic fields, a clinically approved TMS device is capable of activating transduced neurons and eliciting motor freezing in Nb-Ft-TRPV1^Ca2+^ expressing mice at well below the maximal output of the device.

**Fig. 6.**
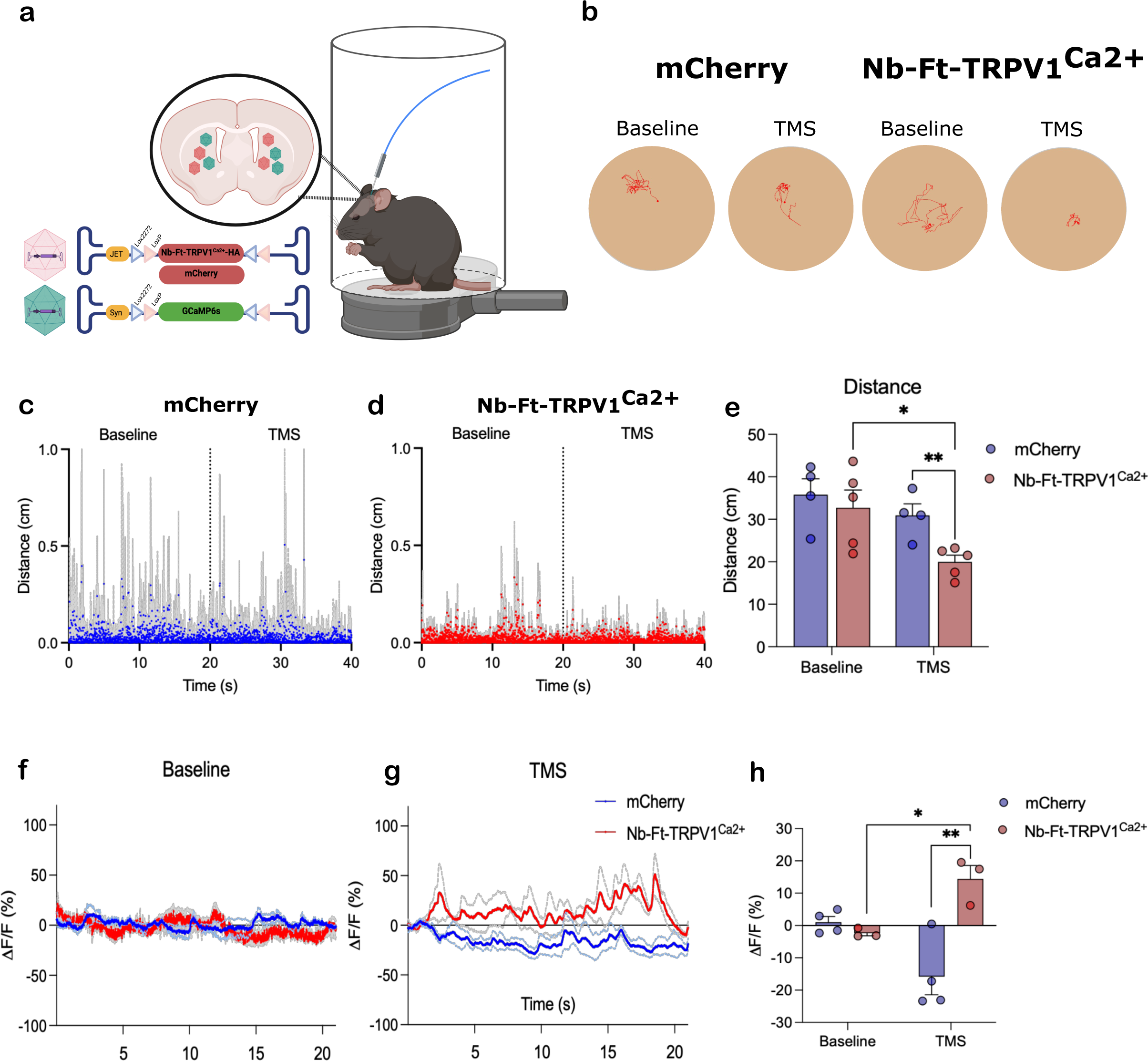
Transcranial magnetic stimulation (TMS) treatment increases Nb-Ft-TRPV1^Ca2+^ mediated motor freezing and calcium transients in dorsal striatum. **A.** Schema of the TMS set up and A2a-Cre mice with bilateral injection of the double-floxed Cre-dependent AAV vector expressing the Nb-Ft-TRPV1^Ca2+^ or mCherry under the control of the JET promoter (red) combined with double floxed Cre-dependent AAV vector expressing GCaMP6s under the control of the Syn promoter (green) in dorsal striatum. **B.** Activity trace example of altered motor activity during baseline (left) and TMS (right) treatment in an mCherry mouse (blue), and Nb-Ft-TRPV1^Ca2+^ mouse (red). Distance ambulated during baseline (left) and TMS (right) treatment in the **C.** mCherry group (blue), and **D.** Nb-Ft-TRPV1^Ca2+^ group (red). (n=4-5; data presented as the mean ± SEM in grey.). **E.** Average distance during baseline and TMS treatment in mCherry (blue) and Nb-Ft-TRPV1^Ca2+^ groups. **F.** Mean fluorescence during baseline in Nb-Ft-TRPV1^Ca2+^ (red) and mCherry (blue) mice (n=3; data presented as the mean ± SEM). **G.** Mean fluorescence during 20 seconds of TMS triggers every 2 seconds in Nb-Ft-TRPV1^Ca2+^ (red) and mCherry (blue) mice (n=3-4; data presented as the mean ± SEM). **H.** Grouped bar graph of Nb-Ft-TRPV1^Ca2+^ (red) and mCherry (blue) mice during baseline, and 20 seconds of TMS triggers every 2 seconds (n=3-4). Error bars show SEM. ** represents p value < 0.01, *** represents p value < 0.001 with two-tailed, unpaired (or paired when baseline vs TMS comparison were done) t-test with Welch’s correction.

### Magnetogenetic stimulation of striatopallidal neurons in wild type mice reproduces a robust motor phenotype

Cre-driver mouse lines provide cell specificity for functional mapping of neural circuits underlying numerous behaviors and many diseases, but this approach is limited to animals. In addition, expression is based on cellular identity and not on functional anatomical connections (*21*). In order to determine if magnetogenetics could be restricted to specific circuits in normal mice, we utilized a dual vector system to restrict expression to the striatopallidal iSPN circuitry. AAV1/2-JET-DIO-Nb-Ft-TRPV1^Ca2+^-HA was first injected bilaterally into the dorsal striatum of normal mice, which would not lead to expression of the viral transgene due to the absence of cre. This was followed by a second injection of a retrograde AAV vector carrying a cre recombinase fused to GFP (RetroAAV2-CMV-Cre-GFP) into the globus pallidus (Gp). This approach allows retrograde uptake of AAV into afferents to the Gp, with resulting cre expression in neurons which project to Gp, including D2 iSPNs. Since the only pallidal afferents containing a floxed transgene with this approach would be those from the dorsal striatum, this should lead to restricted expression of the magnetogenetic construct only in striatopallidal neurons. As expected, immunostaining for HA or RFP (for AAV1/2-JET-DIO and the mCherry control group, respectively) demonstrated co-localization of each with GFP from the transduced SPNs **(Fig. 7a)**. The same behavioral tests were then performed as were used to previously characterize the transgenic A2a-cre mice (**Fig.2**). Consistent with the prior results, there was a robust decrease in locomotor activity when the animals receiving the AAV1/2-JET-DIO-Nb-Ft-TRPV1^Ca2+^-HA injections were placed in a magnetic field compared to controls (**Fig. 7b)**. There was a significant decrease in ambulation during DMF application (n=8, Nb-Ft-TRPV1^Ca2+^ + DMF= 16.13 ± 3.02 vs mCherry + DMF= 39.38 ± 3.28 sec, p<0.001), and significant increase in time spent freezing (n=8, Nb-Ft-TRPV1^Ca2+^ + DMF= 23.75 ± 4.63 vs mCherry + DMF= 4.13 ± 1.34 sec, p<0.001) **(Fig. 7c & d)**. *c-fos* expression was again significantly higher in the transduced iSPNs of Nb-Ft-TRPV1^Ca2+^ expressing mice compared to mCherry controls (n=4, Nb-Ft-TRPV1^Ca2+^ + DMF= 26.34 ± 3.13 vs mCherry + DMF= 10.76 ± 2.41 %, p<0.01) **(Fig. 7e & f).** To control for the possibility that striatal overexpression of TRPV1 ion channels alone could mediate activation of the indirect pathway in the absence of ferritin tethered to the TRPV1 channel, we compared the effects of DMF treatment on wt mice expressing either Nb-Ft-TRPV1^Ca2+^ or TRPV1^Ca2+^ alone (**Fig. 7g)**. DMF application had no observable motor effect on TRPV1^Ca2+^ expressing mice but again suppressed ambulation and increased freezing in mice expressing Nb-Ft-TRPV1^Ca2+^, confirming that fusion of the ferritin nanobody to the TRPV1 channel was required for neuronal activation in the presence of a magnetic field **(Fig. 7h & i)**.

**Fig. 7.**
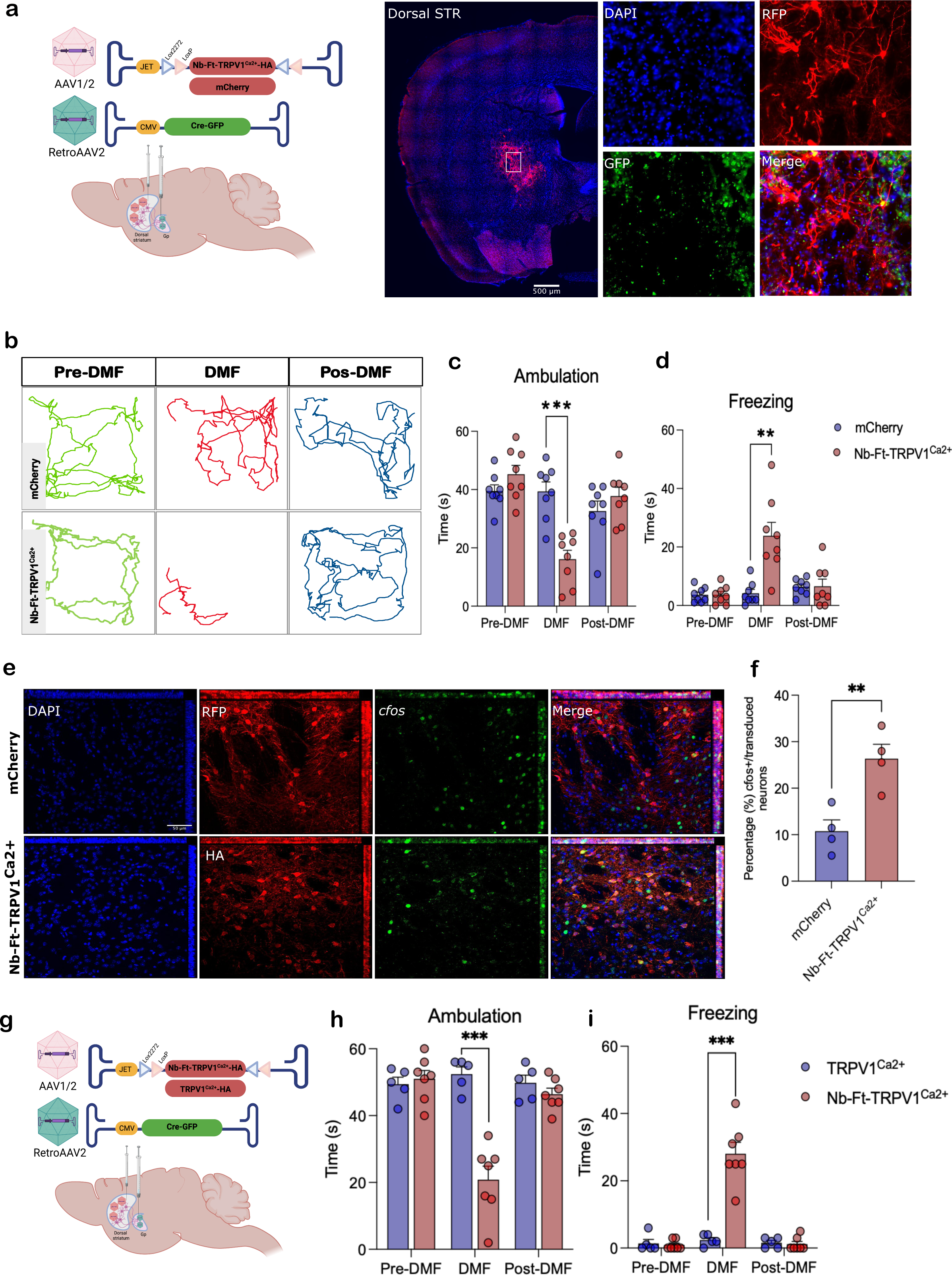
Selective viral-mediated expression of Nb-Ft-TRPV1^Ca2+^ in the striatopallidal pathway elicit parkinsonian motor behavior in wild type mice. **A.** Schema of the double-floxed Cre-dependent AAV vector expressing the Nb-Ft-TRPV1^Ca2+^ or mCherry under the control of the JET promoter in striatal iSPN projecting to the Gp achieved by Gp injection of a retrograde AAV2 vector carrying Cre-GFP, and immunostaining for GFP (green), HA (red) in wt mice demonstrates selective expression of the AAV-Nb-Ft-TRPV1^Ca2+^ in dorsal striatum iSPN projecting to the Gpe, white arrows show colocalization of GFP with HA. **B.** Example of altered motor activity during bilateral striatopallidal pre-DMF (green), DMF (red) and post-DMF (blue) stimulation, lines represent the mouse’s path. Effect of DMF stimulation on **C.** Ambulation bout duration, and **D.** Freezing of gait in mCherry (blue bars/dots, n=8) and Nb-Ft-TRPV1^Ca2+^ (red bars/dots, n=8) mice. **E.** Immunostaining for *c-fos* (cyan) and HA (Nb-Ft-TRVP1^Ca2+^; red) or RFP (mCherry; red) of dorsal striatum of wt mice euthanized 1 hour following 15 minutes of DMF exposure. White arrows show co-localization. **F.** Percentage (%) of *c-fos* positive (+) neurons over total number of iSPNs expressing HA or RFP (n=3 mice per group/3 sections per mice). **G.** Schema of the double-floxed Cre-dependent AAV vector expressing the Nb-Ft-TRPV1^Ca2+^ or TRPV1^Ca2+^ under the control of the JET promoter in striatal iSPNs projecting to the Gp achieved by Gp injection of a retrograde AAV2 vector carrying Cre-GFP. Effect of DMF stimulation on **H.** Ambulation bout duration, and **I.** Freezing of gait in TRPV1^Ca2+^ (blue bars/dots, n=5) and Nb-Ft-TRPV1^Ca2+^ (red bars/dots, n=7) mice. Error bars show SEM. ** represents p value < 0.01, *** represents p value < 0.001 with two-tailed, unpaired t-test with Welch’s correction .

### Magnetogenetic inhibition of subthalamic nucleus neurons improves motor impairment in parkinsonian mice

One advantage of chemogenetic and optogenetic systems is the availability of a suite of modified channels which can inhibit neurons in addition to those which can activate neuronal firing (*22*). Developing an inhibitory magnetogenetic system would not only expand the opportunities for scientific research using this technology but would also provide translational opportunities for diseases where neuronal inhibition would be desirable. We first validated *in vitro* the utility of the anti-ferritin nanobody fused to a mutant TRPV1 channel modified to allow permeability of Chloride (Cl-) ions (Nb-Ft-TRPV1^Cl-^), which should increase chloride flux to inhibit neuronal activity **(Fig. 8a)**. Neuro2A cells were transfected with plasmids expressing mCherry, TRPV1^Ca2+^ or Nb-Ft-TRPV1^Cl-^ for fluorescent chloride imaging using MQAE, which is quenched by intracellular chloride **(Supplementary Fig. 8a)**. Application of a low intensity magnetic field (MF, 230-250mT) showed a significant decrease in the ΔF/F0 in the Nb-Ft-TRPV1^Cl-^ expressing cells compared to both mCherry and TRPV1^Ca2+^ controls **(Fig. 8b & c)** (ΔF/F0 at 150s: Nb-Ft-TRPV^Cl-^-0.16 ± 0.01, TRPV1^Ca2+^0.06 ± 0.01, mCherry -0.002 ± 0.01, p<0.001) consistent with increased intracellular chloride concentrations.

**Fig. 8.**
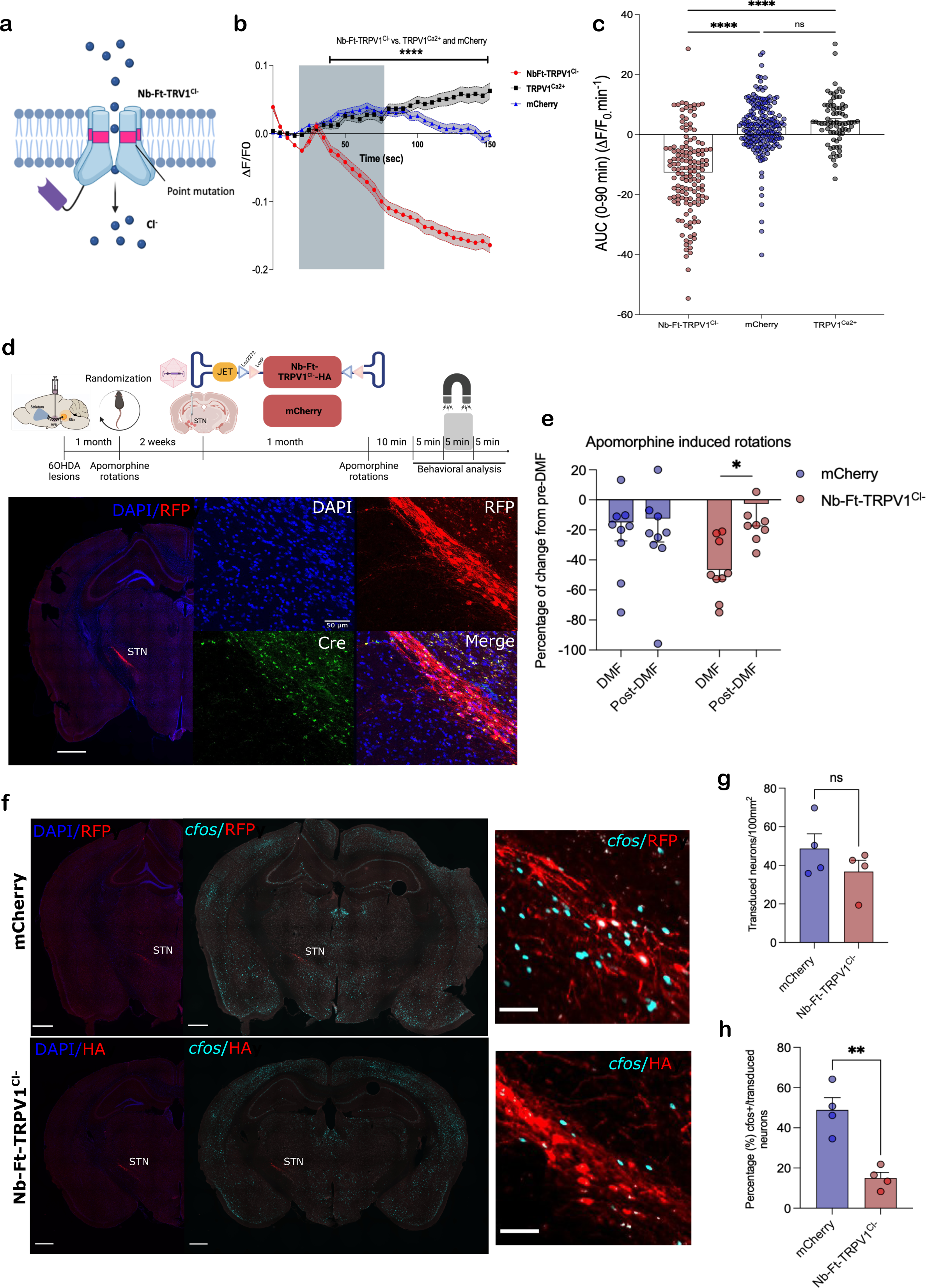
Mutant Nb-Ft-TRPV1^Cl-^ inhibits neuronal activity of subthalamic nucleus projection neurons and rescues motor impairment in parkinsonian mice.A. Schema of inhibition system with mutant Nb-Ft-TRPV1^Cl-^ membrane channel. **B.** Normalized change in MQAE fluorescence Intensity (ΔF/F0) of Neuro2A cells expressing mCherry, TRPV1^Ca2+^ or Nb-Ft-TRPV1^Cl-^ with low AMF application *in vitro*. **C.** AUC analysis of chloride imaging for Nb-Ft-TRPV1^Cl-^ channel. **D.** Protocol and schema for generation of parkinsonian PitX-2-Cre mice with unilateral injection of 6 hydroxydopamine (6-OHDA) in the medial forebrain bundle (MFB), followed by randomization based contralateral rotations induced by i.p. administration of apomorphine (0.25mg/kg) and ipsilateral intracranial injection of the double-floxed Cre-dependent AAV vector expressing the mutant Nb-Ft-TRPV1^Cl-^ under the control of the JET promoter in the subthalamic nucleus (STN). Representative immunostaining of the selective expression of Nb-Ft-TRPV1^Cl-^ (red) in STN neurons expressing Cre recombinase (Green). **E.** Percentage of change from pre-DMF treatment in contralateral rotations induced with apomorphine (0.25mg/kg, i.p.) in parkinsonian PitX-2-Cre mice expressing the Nb-Ft-TRPV1^Cl-^ or the mCherry control viruses in the STN during DMF, and post-DMF. **F.** Immunostaining for *c-fos* (cyan) and HA or mCherry (red) of STN of PiX-2-Cre mice euthanized 1 hour following 15 minutes of DMF exposure with apomorphine induced rotations (0.25mg/kg).**G**. Number of transduced neurons per area in STN between mCherry and Nb-Ft-TRPV1^Ca2+^ groups. **H.** Percentage (%) of *c-fos* positive (+) neurons over total number of STN neurons expressing HA or RFP (n=4 mice per group/3 sections per mice). Error bars show SEM. * represents p value < 0.05, ** represents p value < 0.01 with two-tailed, unpaired t-test with Welch’s correction. Anti-HA staining to detect Nb-Ft-TRVP1^Ca2+^ expression, anti-RFP staining to detect mCherry expression.

The use of an inhibitory magnetogenetic system may be of potential use for symptomatic treatment of Parkinson’s disease (PD), where chemical or gene therapy-based inhibition of subthalamic nucleus (STN) projection neurons can improve motor function in rodents, non-human primates, and humans (*23–27*). To test this, we generated unilateral 6-hydroxydopamine (6-OHDA) lesions of nigral dopaminergic neurons in transgenic mice expressing the cre recombinase in PitX-2-positive neurons (PitX-2-cre mice) **(Supplementary Fig. 8b)**, which restricts cre expression to STN glutamatergic projection neurons with virtually no expression within other populations in the vicinity (*28*). Stable and functionally significant lesions were documented through quantification of apomorphine-induced rotations, followed by randomization of animals to balanced experimental and control groups and confirmation that DMF alone does not affect rotations **(Supplementary Fig. 8c-d)**. A DIO cre-dependent AAV1/2 vector encoding the Nb-Ft-TRPV1^Cl-^ cassette under the control of the JET promoter (AAV1/2-JET-DIO-Nb-Ft-TRPV1^Cl-^-HA) was then injected into the ipsilateral STN of 6OHDA-lesioned PitX2 mice, with another group injected with a DIO cre-dependent mCherry vector as a negative control **(Fig. 8d)**. Mice expressing Nb-Ft-TRPV1^Cl-^ in the STN showed a significant reduction of contralateral rotations induced by the dopaminergic agonist, apomorphine, in the presence of a DMF application relative to baseline and controls baseline (n=9-10, -46.62 ± 6.5 vs -2.67 ± 14.69%, p<0.05), **(Fig. 8e, Supplementary Fig. 8e & f, supplementary data file)**. The number of *c-fos* positive transduced neurons in the STN of PitX2-cre mice was significantly decreased following DMF exposure in the group which received the inhibitory magnetogenetic Nb-Ft-TRPV1^Cl-^vector compared to mCherry controls **(Fig. 8f & g)** (n=4, Nb-Ft-TRPV1^Cl-^= 15.01 ± 2.83 vs mCherry= 48.89 ± 6.11%, p<0.01). This confirms that AAV-mediated expression of the magnetogenetic inhibitory construct exclusively within PitX2-positive STN projection neurons resulted in reduction of neuronal activity *in vivo* and a consequent improvement in motor function following 6OHDA lesioning of the substantia nigra.

## Discussion

The demonstration that viral vectors such as AAV can be used for gene transfer in the mammalian brain (*29*) has transformed pre-clinical neuroscience research. Furthermore, many completed and active clinical trials of gene therapy in humans have shown promise for treating circuit disorders (*27, 30*). While initially genes cloned into AAV were expressed constitutively, more recently techniques such as optogenetics and chemogenetics have been used to regulate neural activity in an inducible manner, using light or drugs respectively. Optogenetics provides temporally rapid and precise control of neuronal function but utilizes microbial-derived channels that can be immunogenic and require permanent implantation of light-generating devices that potentially tether small animals and present engineering and safety challenges for human applications (*4, 31*). Chemogenetics uses drugs to regulate channels, which obviates the need for a device implant, but chemogenetic regulation occurs in a more delayed fashion based upon the pharmacokinetics of the activator molecules (*32, 33*). In this report, we validate magnetogenetics as an alternative approach for modulating neural activity and show that magnetic field-based regulation of neuronal function can be used for bidirectional control of neuronal activity. This method provides both anatomical and temporal precision in awake, freely moving animals without the need for an implanted device and uses a viral vector potentially applicable for clinical studies and can respond to clinically approved devices which deliver sufficient strength magnetic fields.

We created this system using variants of TRPV1, which are endogenous mammalian channels, together with a nano-body that binds to endogenous mouse and human ferritin(*34*). TRPV1 is attractive as a channel as it is less sensitive to temperature and osmotic changes than some other TRP channels such as TRPV4, which may reduce non-specific activation of the channel itself in the absence of a magnetic field(*35*). We tested this approach by eliciting a parkinsonian phenotype in freely moving mice to demonstrate the functional efficacy of magnetogenetic activation of striatal indirect medium spinal neurons. Activation of these neurons in a magnetic field was evident only in mice receiving the magnetogenetic construct compared with controls, as confirmed by *c-fos* expression post-mortem as well as both FDG-PET and fiber photometry in living animals. Optogenetic control of basal ganglia circuitry has previously demonstrated similar changes in motor behavior after selective activation of indirect pathways *in vivo* using an implanted light device to control neurons expressing Channelrhodopsin-2 (ChR2) (*14*). More recently, Munshi and colleagues (*5*) reported successful magnetogenetic modulation of motor circuits targeting motor cortex and dorsal striatum. Consistent with the classical model of basal ganglia circuitry, this study reported decrease in voluntary movement by activation of striatal D2 iSPN using a combination of metal nanoparticles (MNPs) and AAVs encoding TRPV1 channels stereotactically injected 4 weeks apart in the ridge between dorsal and ventral striatum (*5*). However, synthetic magnetic nanoparticles would not allow for stable, chronic magnetogenetic activation and the use of MNPs could potentially tether to other thermo-sensitive channels causing off target activation and safety issues (*7, 36*). In addition, the intrinsic variability of separate injections and non-cell specific transgene transduction can limit conclusions regarding the relationship between phenotype and cells modulated by the exogenous particle-based magnetogenetic system. This is in contrast to the genetically encoded system which we employed, which only requires injection of a single AAV to improve both dosing consistency between subjects and longevity of neuromodulation due to chronic AAV-mediated expression. For human translation, the ability to deliver a single construct with a single infusion has both safety and regulatory advantages, and here we demonstrate the viability of AAV-mediated delivery of a single magnetogenetic construct encoding a fused Nb capable of tethering endogenous ferritin to an excitatory TRPV1 channel for inducible modulation of neuronal activity leading to definable motor phenotypes.

The ability use of optogenetics, through a combination of focal delivery of viral constructs expressing opsins and implantation of light emitting devices in a target brain region, has led to a wealth of knowledge regarding the relevance of neuronal circuits to particular behaviors (*37*). The potential benefit of the magnetogenetic system described here is the use of a non-invasive magnetic field to regulate specific circuits in freely moving animals, rather than a permanent optical fiber implant which is tethered to an external device. We first demonstrated that a cre-dependent magnetogenetic system can target particular neuronal populations in cre-driver mice. However, this approach is limited to animals and may also not provide the anatomical specificity required to examine anatomical circuits. To address this, we also developed a modified approach combining delivery of the cre-dependent magnetogenetic activating construct into the striatum with injection of a retrograde viral vector (AAVretro) into globus pallidus, a known target of striatal iSPNs, to restrict expression of the activating construct exclusively within the same striatopallidal iSPNs targeted in the A2a-cre driver mice. Histological analysis confirmed that expression was restricted to striatal D2 neurons and the behavioral and physiological responses to the magnetic field were similarly robust compared with data generated in the A2a-cre mice. However, AAVretro may not efficiently transduce all neurons in a retrograde fashion and must be individually tested for particular applications, though there are alternative retrograde viral vectors that could also be used in such an approach if the pattern of retrograde uptake differs between vectors (*38*). Nonetheless, our data provide a novel means for non-invasive regulation of specific neuronal circuits without the need for transgenic animals, which could not only be useful for pre-clinical research but even have potential for circuit-specific human translation.

One possible limitation of the application of magnetogenetics in a clinical setting is the availability of a device for generating a magnetic field that is suitable for human therapeutic applications. Stanley and colleagues (*10*) previously reported successful bidirectional magnetogenetic modulation of ventromedial hypothalamus (VMH) neurons using the earlier dual construct system and an MRI to deliver the magnetic field, and we similarly observed a robust motor phenotype following exposure to a 3T MRI in mice expressing Nb-Ft-TRPV1^Ca2+^ in iSPNs at similar field strength thresholds. While this is feasible for experimental studies, any requirement of proximity to a 3T MRI for efficacy could limit the translational potential of this technology. We addressed this by also testing a TMS device as a source of AMF to induce magnetic-field-based stimulation of D2 iSPNs. As with the MRI machine, we observed a sustained effect from the TMS generated magnetic field in mice expressing Nb-Ft-TRPV1^Ca2+^ in striatal D2 iSPNs, with decreased locomotion and corresponding increase in GCaMP fluorescence. Interestingly, *in vivo* fiber photometry analysis also revealed that TMS reduced neural activity in control mice, which could potentially be attributed to TMS frequency-dependent neural activation-inhibition. This is not surprising, as TMS does influence neuronal activity in isolation, which is the basis for current clinical applications. Human (*39*) and animal (*20*) reports that have shown suppression of neural activity in motor-related regions with different protocols of TMS treatment, especially low-frequency TMS. Nonetheless, this reduced activity had no behavioral effect and if anything suggests that we may even have underestimated the magnitude of magnetogenetic neuronal activation. Given the widespread and increasing use of TMS as a clinical device for non-invasive neuromodulation in a variety of conditions (*19*), it is possible that magnetogenetics could be used to either augment TMS or allow TMS to target deeper neuronal populations where the strength of the magnetic field weakens but is still above the threshold that we have observed for functional activation of the recombinant engineered channel. The ability to generate low field strength at-home devices for intermittent modulation similar to TMS, while maintaining anatomical specificity through focal delivery of the gene therapeutic, could also allow broader application of this technology. Finally, the ability to regulate peripheral or autonomic neurons is plausible using this technology for indications where either activation or inhibition of neural activity is desirable, potentially including the control of pain, neuropathy symptoms or organ function. Our data establishing the threshold for magnetic field strength also raises the possibility that other more simple or even wearable magnetic devices may enable regulation of more accessible neuronal populations.

To expand the utility of this system, we have also developed a novel construct with a mutation that converted the sodium channel into a chloride channel, allowing magnetogenetic inhibition of neuronal activity. Our *in vitro* data demonstrated the functionality of this new construct, while AAV-mediated delivery of the construct to a population of glutamatergic projection neurons within the STN of hemiparkinsonian PitX2-cre mice confirmed the *in vivo* efficacy of magnetogenetic inhibition and consequent motor improvements in this setting. Pitx2 expression is restricted to neurons within the general region of the STN (*40, 41*), allowing very limited expression of our cre-conditional magnetogenetic system to this population of STN neurons. Altered activity of the STN is widely believed to be one of the pathophysiological mechanisms leading to PD motor deficits (*42*), hence traditional high frequency DBS in the STN has emerged as a standard therapy for advance stages of PD (*2*). Nonetheless, there is equipoise regarding the mechanism of DBS action and the role of inhibition of STN neurons as a PD therapeutic. Gradinaru and colleagues (*42*) found that optical inhibition of glutamatergic STN neurons in hemi-parkinsonian rats was insufficient to affect motor behavior using early generation inhibitory opsins, yet STN lesioning, infusion of GABA agonists into the STN and gene therapy to promote GABA production within the STN have all demonstrated improved motor function in both pre-clinical studies and human clinical trials (*23–25, 27, 43*). While DBS may still function through mechanisms other than simple neuronal inhibition (*44*), our results support a significant body of evidence suggesting that inhibition of STN projection neurons can improve motor function in Parkinson’s disease (*23, 27, 45–48*). The availability of an inhibitory magnetogenetic system, which can also be combined with retrograde cre vectors to target specific populations and can be applied to peripheral neurons as well, could vastly expand both the experimental and clinical potential of this technology.

The data reported here firmly establish that expression of a TRPV1 channel tethered to ferritin *in vivo* regulates neuronal activity in the presence of a magnetic field using several different independent readouts, including real-time calcium signaling, PET-based glucose utilization and immediate early gene expression. Together with the corresponding behavioral changes, these data confirm that the TRPV1-ferritin fusion proteins that we developed can bidirectionally modulate neural activity *in vivo* in response to a magnetic field of sufficient strength. While other groups have also published empiric data confirming a cellular response with magnetogenetic activating constructs using different readouts(*7, 49*), the possibility that mechanical or thermal gating as a mechanism of regulating the channel was initially questioned on purely theoretical grounds(*50*). However, more recently several mechanisms have been proposed to explain gating of TRP channels tethered to ferritin in a magnetic field. Detailed theoretical analyses of a range of potential physical parameters of ferritin-iron nanoparticles and their interaction with the mechanosensitive ion channel suggest the possibility that alternative physical forces can trigger channel opening(*51*). These include possible magneto-caloric and magneto-thermal mechanisms, with another study raising the possibility that a magneto-caloric mechanism induces a thermally mediated magnetic response via magnetic entropy(*52*). In addition, several groups have generated data to support a possible chemical mechanism, in which exposure of ferritin to either a magnetic or radiofrequency field increases the cellular levels of reactive oxygen species with lipid oxidation changing TRP channel dynamics(*36, 53, 54*). These possibilities are not mutually exclusive and both chemical and physical factors may contribute. Bidirectional control of neuronal activity through the unimolecular fusion proteins described here which control the relationship between the TRPV1 mutants and the ferritin nanobody should also provide advantageous tools to further analyze these possible mechanisms.

In summary, the present study demonstrates that magnetogenetic gene therapy efficiently regulates basal ganglia circuitry. A single AAV encoding the activating magnetogenetic construct restricted to iSPNs elicited a freezing phenotype in transgenic mice. Circuit-specific magnetogenetic activation of iSPNs was replicated in wt mice using AAVretro to deliver cre to a specific neuronal population, while expression of a novel inhibitory magnetogenetic construct within the STN reduced activity with consequent motor improvement in hemiparkinsonian mice. Our results provide a bidirectional toolkit for magnetogenetic regulation to study specific central or peripheral neural circuits, while further development of the technology may facilitate translation into amenable human conditions where non-invasive and immediate modulation of neuronal activity could have therapeutic benefit.

## Materials and Methods

### Ferritin nanobody generation

Immunization of a llama: A llama was injected subcutaneously with human spleen ferritin (250 μg) (Lee Biosolutions, MO) on days 0, 7, 14, 21, 28 and 35. Gerbu LQ#3000 was used as adjuvant. On day 40, anticoagulated blood was collected for lymphocyte preparation.

#### Construction of a VHH library

A VHH library was constructed and screened for the presence of antigen specific nanobodies. To this end, total RNA from peripheral blood lymphocytes was used as template for first strand cDNA synthesis with oligo(dT) primer. Using this cDNA, the VHH encoding sequences were amplified by PCR, digested with PstI and NotI, and cloned into the PstI and NotI sites of the phagemid vector pMECS. A VHH library of about 108 independent transformants was obtained. About 87% of transformants harbored the vector with the right insert size.

#### Isolation of antigen-specific nanobodies

The library was subject to three consecutive rounds of panning on solid-phase coated human ferritin (200 µg/ml, 20 μg/well). The enrichment for human ferritin-specific phages was assessed after each round of panning by comparing the number of phagemid particles eluted from antigen coated wells with the number of phagemid particles eluted from negative control wells blocked without antigen. These experiments suggested that the phage population was enriched about 20-fold, 20-fold, and 8 × 102 -fold for human ferritin-specific phages after 1st, 2nd and 3rd rounds of panning, respectively. 190 colonies from 1st & 2nd rounds were randomly selected and analyzed by ELISA for the presence of human ferritin-specific nanobodies. This first round of ELISAs was used to screen nanobodies in crude periplasmic extracts and the relative intensity of the ELISA signal may not reflect the relative quality of the nanobodies. In these experiments the differences in ELISA signals between different nanobodies may be related to variables such as the amount of nanobody, rather than to nanobody quality such as affinity or actual yield. Out of these 190 colonies, 133 colonies scored positive for the presence of nanobodies to human ferritin in this assay (46 & 87 from 1st & 2nd rounds, respectively). The human ferritin used for panning and ELISA screening was the same as the one used for immunization. Based on sequence data, the ELISA-positive colonies represented 59 different nanobodies belong to 22 different CDR3 groups.

Next, the library was subject to three consecutive rounds of panning on solid phase coated with mouse ferritin purified from mouse liver (200 µg/ml, 20 μg/well). The enrichment for mouse ferritin-specific phages was assessed as above. These experiments suggested that the phage population was enriched about 10-fold, 10- fold and 5 × 102 -fold for mouse ferritin-specific phages after 1st, 2nd, and 3rd rounds of panning, respectively. 190 colonies from 2nd & 3rd rounds were randomly selected and analyzed by ELISA for the presence of mouse ferritin-specific nanobodies in their periplasmic extracts (ELISA using crude periplasmic extracts including soluble nanobodies). Out of these 190 colonies, 54 colonies scored positive in this assay (1 & 53 from 2nd & 3rd rounds, respectively). Based on sequence data, the ELISA-positive colonies represented 19 different nanobodies belonging to 6 different CDR3 groups. Two nanobody CDR3 groups bound both human and mouse ferritin.

### Purification of recombinant nanobody protein

pMECS vectors encoding anti-ferritin nanobodies were transformed into electrocompetent E.coli WK6 cells. Colonies were picked and grown in 10-20 ml of LB + ampicillin (100 µg/ml) + glucose (1%) at 37°C overnight. 1ml of preculture was added to 330 ml TB-medium (2.3 g KH2PO4, 16.4 g K2HPO4.3H2O,12 g Tryptone, 24 g Yeast and 4 ml 100% glycerol supplemented with 100 µg/ml Ampicillin, 2mM MgCl2 and 0.1% glucose and grow at 37°C. Nanobody expression was induced by addition of IPTG (1mM). The culture was incubated at 28°C with shaking overnight. The cells were pelleted for 8 minutes at 8000 rpm and resuspended in 12 ml TES (0.2 M Tris pH8.0, 0.5 mM EDTA, 0.5 M sucrose) before shaking for 1 hour on ice. A further 18ml of TES (diluted 4-fold in water) was added and incubated for a further hour on ice with shaking then centrifuged for 30 mins at 8000rpm, at 4°C. The supernatant containing the nanobody proteins was then purified by immobilized metal-ion affinity chromatography (IMAC) using HIS-Select® Cobalt Affinity Gel (Sigma H8162) according to the manufacturer’s instructions. The amount of protein was estimated at this point by OD280 measurement of eluted sample. 14 nanobodies were successfully purified.

### Characterization of nanobody affinity to human ferritin by ELISA

ELISA plates were coated with 1µg/mL human spleen ferritin (Lee Biosolutions, MO) in PBS overnight at 4°C. Plates were then washed 5 times with washing buffer (PBS+0.05% Tween-20), followed by blocking with PBS+ 1% BSA at room temperature for 2 hrs. Serial dilutions of nanobodies or BSA (100ul) were added to the ELISA plates and incubated for 2 hours at room temperature then washed 5 times with buffer. Anti-HA-HRP antibody (100ul, Miltenyi biotech, 130-091-972, 1:1000 dilution) was added to each well and incubated for 1 hr at RT before a further 5 washes. TMB substrate (100ul) was added to each well and the reaction stopped by addition of 100 µL 2M H2SO4 after 10 mins. OD450 was measured using a microplate reader (BMG Labtech CLARIOstar plate reader).

### Characterization of anti-ferritin nanobodies by immunoprecipitation

Nanobody sequences were amplified by PCR with the specific primers (NbFt-F: gcaagatctgccaccatggcc CAGGTGCAGCTGCAGGAG; NbFt-R:gcaaagcttggatccAGCGTAATCTGGAACATCGTATGGGTA tgcggccgctgagga) and subcloned into the retroviral vector pMSCV (Clontech, Takara Bio, US). HEK-293T cells were grown in 10cm plates and co-transfected with plasmids expressing GFP-mFerritin and anti-ferritin nanobody clones using PEI (DNA:PEI=1:3). Cells were harvested after 48 hours and lysed with 500 µl of lysis buffer (50 mM Tris, pH 7.5, 300 mM NaCl, 1 mM EGTA, 1 mM EDTA, 1% NP-40, 0.1% SDS) and protease inhibitor (complete EDTA-free, Roche) then incubated for 15 mins with agitation at 4o C before centrifugation at 14,000 rpm for 20 mins. The supernatant was transferred to a new tube with 30 µl HA.11 antibody conjugated agarose beads (Biolegend, 900801) added to each sample. The beads and supernatant were incubated at 4o C overnight with rotation and then spun at 3000rpm for 2 mins. Beads were then washed 3 times with lysis buffer before proteins were eluted with 20 µl of 4x Laemmili buffer and heated at 95o C for 5 minutes. Eluted proteins were examined by SDS-PAGE. Samples were loaded onto a 10 well SDS-PAGE gel and run for 10-15mm then fixed in 46% methanol/7% glacial acetic acid for 1 hour. The gel was then stained for 1 hour in 0.1% Coomassie blue R-250 in 46% methanol/ 7% glacial acetic acid before destaining in ultrapure water. Bands were then excised and analyzed by mass spectrometry.

### Mass spectrometry

Proteins were reduced (10 mM DTT, EMD Millipore) and alkylated (30 mM iodoacetamide, Sigma), followed by digestion with Endoproteinase LysC (Wako Chemicals) and trypsin (Promega). Reaction was halted by addition of trifluoracetic acid and peptides were solid phase extracted (46) and analyzed by reversed phase nano-LC-MS/MS (Dionex 3000 coupled to a Q-Excative Plus, Thermo Scientific). Data were quantified and searched against a Uniprot human database concatenated with a mouse GFP-Ferritin sequence using ProteomeDiscoverer 1.4/Mascot 2.5. Oxidation of methionine and protein N-terminal acetylation were allowed as variable modifications. Cysteines were considered as fully carbamidomethylated. Two missed cleavages were allowed. Peptide matches were filtered using a percolator(47) calculated false discovery rate of 1%. The average area of the three most abundant peptides per protein were used as a proxy for protein abundance (48). For the highly homologous ferritin heavy proteins, only peptides unique to either the mouse or the human forms were used in access abundance. Peptides specific to the N-terminal GFP moiety of the over expressed mouse ferritin was not used to calculate abundance of mouse ferritin but were recorded separately.

### Isothermal titration calorimetry

We performed an isothermal titration calorimetry (ITC) assay using a MicroCal Auto-iTC200 instrument (Malvern Panalytical). In each experiment, 10 μM human spleen ferritin in the cell was titrated with 75 μM NbFt2 in the syringe at 25 °C. The titration sequences included a single 0.4 μL injection, followed by 19 injections of 2 μL each, with a 150-sec interval between injections, stirring rate of 750 rpm and reference power of 10 μcal/sec. Data were analyzed with Origin analysis software.

### Generation of constructs

The expression vectors for calcium dependent release of insulin and SEAP, pCMV-TRPV1Ca2+ were generated as previously described(20, 21). To generate Ca2+-dependent SEAP reporter, the SEAP coding fragments were obtained by cutting pYSEAP (addgene 37326) with HindIII and HpaI and inserted into pSRE-CRE-NFATinsulin(21) at the HindIII and HpaI sites. To express Nb-FT in mammalian cells, a PGK promoter-driven mCherry expression cassette and the WPRE element were inserted into pMSCV (Clontech, Takara Bio) at the XhoI and ClaI sites to generate an empty vector (pMSCV-mCherry-WPRE). The anti-ferritin nanobody coding sequences were amplified from phage display plasmids with specific primers: 5’gcaagatctgccaccatggccCAGGTGCAGCTGCAGGAG and 5’gcaaagctt ggatccAGCGTAATCTGGAACATCGTATGGGTAtgcggccgctgagga, digested with BglII and HindIII and ligated into pJFXY21 at the BglII and HindIII sites. The resulting plasmids were named pMSCV-Nb-Ft-2, -9, -10, -14 and -17-TRPV1Ca2+. The Nb-GFP fragment was amplified from MSCV-αGFP-TRPV1-2AGFPferritin(16) with specific primers: 5’gcaagatctgccaccatggcc CAGGTGCAGCTGCAGGAG and 5’ tcagacgtcggccactgcggccgcTGAGGAGACGGTGACCTGGGTC, digested and subcloned into pNb-Ft-2-TRPV1Ca2+ at the BglII and NotI sites to generate plasmid pNb-GFP-TRPV1Ca2+. The pMSCV-Nb-GFP-TRPV1Ca2+-T2A-GFP-mFerritin plasmid was generated by subcloning the Nb-GFPTRPV1Ca2+-T2A-GFP-mFerritin fragment cut from MSCV-αGFP-TRPV1-2A-GFPferritin(16) with NotI and EcoRI into a modified pMSCV-WPRE vector at the NotI and MfeI. NbFt sequences were amplified from plasmids pMSCV-Nb-Ft-2, -9, -10, -14 and -17-TRPV1Ca2+ with specific primers: 5’ gtgatgcatGCCACCATGGCCCAGGTGCAG and 5’ gatgctagcgccAGCGTAATCTGGAACATCG, digested with NsiI and Nhel and inserted into pJFXY16 at the NsiI and AvrII sites. The resulting constructs were named pMSCV-Nb-Ft-2, -9, - 10, -14 and -17-TRPV1Ca2+-T2A-GFP-mFerritin. The pMSCV-GFP-mFerritin vector was created by digesting pMSCV-Nb-GFP-TRPV1Ca2+-T2A-GFP-mFerritin with BglII to remove the NbGFP-TRPV1-T2A fragment and self-ligated.

To make pAAV-hSyn-Nb-FT-2-TRPV1Ca2+, the 2981-bp PCR product containing Nb-FT-2-TRPV1Ca2+was amplified from pMSCV-Nb-Ft-2-TRPV1Ca2+-T2A-GFP-mFerritin. using primers 5’AGCGCAGTCGAGAAGGTACCGGATCCCCCGGTCGCCACCACTAGTATGGCCCAGGTGCAGCTGC and 5’-TTATCGATAAGCTTGATATCGAATTCTTACTTCTCCCCTGGGACCATG, digesting with BamHI and EcoRI and cloning into BamHI/EcoRI-digested pAAV-hSyn-mCherry (Jimenez-Gonzalez et al, 2021, NBME).

To make pAAV-hSyn-RCaMP, a 1441bp PCR product containing jRGECO1a, a red fluorescent calcium sensor protein (RCaMP) was amplified from pAAV.Syn.Flex.NES-jRGECO1a.WPRE.SV40 (Addgene cat#100853) using primers 5’-ACTAGTATGCTGCAGAACGAGCTTGC and ACCGGTCTACTAGTCTCAATTGTCACTTCGCTGTCATCATTTGT, digested with SpeI, and cloned into pAAV-hSyn-DIO-MCS in the forward direction so that RCaMP could be expressed in the absence of Cre recombinase. pAAV-hSyn-DIO-MCS is pAAV-hSyn-DIO-hM3D(Gq)-mCherry (Addgene cat# 44361) which had previously had the BsrGI/NheI region containing hM3D(Gq)-mCherry replaced with the following multiple cloning site: TGTACAACTAGTACCGGTTCGCGAGCATGCCCTAGGGCTAGC.

### Cell culture and in vitro studies

Human embryonic kidney cells (HEK-293T, (ATCC^®^ CRL-3216^™^) mycoplasma testing and STR profiling performed by ATCC) were cultured in Dulbecco’s modified eagle medium with 10% fetal bovine serum (Gibco, Carlsbad, CA) at 37°C and 5% CO_2_. Neuro 2A cells (ATCC CCL-131, mycoplasma testing and STR profiling performed by ATCC) were grown in Eagles’ minimum essential medium with 10% fetal bovine serum (Gibco) at 37°C and 5% CO_2_.

#### Immunocytochemistry studies

HEK-293T or Neuro 2A cells were cultured on 12-mm cover glass (Fisher Scientific, Pittsburgh, PA) coated with fibronectin (Sigma Aldrich, cat#F1141). Cells were transfected with constructs expressing pAAV-hSyn-Nb-FT-2-TRPV1^Ca2+^ or pCMV-TRPV1^Ca2+^ 24 h after plating using X-tremeGENE^™^ 9 DNA Transfection Reagent (Millipore Sigma) according to manufacturer’s specifications. Cells were stained 48-72hrs after transfection.

#### Magnet treatment studies using calcium-dependent reporters

HEK-293T cells were cultured on 12-mm cover glass and transfected with pMSCV-Nb-Ft-TRPV1^Ca2+^ or pMSCV-Nb-GFP-TRPV1 and GFP-mFerritin and either calcium-dependent SEAP construct(*55*) or calcium-dependent insulin construct(*8*). Holotransferrin (2 mg/ml, Sigma) was added to cells 24 hours after transfection. 24 h prior to the study, cells were placed in 1% FBS medium at 32°C to ensure minimal activation of TRPV1 and calcium dependent pathways. Cells were incubated in 300 µl of calcium imaging buffer at room temperature (control) or in an oscillating magnetic field (465kHz, 30mT) at room temperature. After 60 mins, the supernatant was removed, spun to remove cells, and assayed for secreted alkaline phosphatase or insulin. Screening studies with SEAP production were repeated twice with 4 replicates. Validation studies with calcium dependent insulin production were repeated at least 3 times with at least 3 replicates.

#### Calcium imaging studies

Neuro2A cells were cultured on cover glass and transfected with pAAV-hSyn-Nb-FT-2-TRPV1^Ca2+^ with or without pMSCV-GFP-mFerritin as described above with holotransferrin added 24 hours after transfection. Cells were placed at 32°C 24 hours before testing. Cells were loaded with Fluo-4 3µM (Invitrogen) in the presence of sulfinpyrazone 500 µM (Sigma) for 45-60 min at 32°C then washed and incubated for 15-30 min in sulfinpyrazone in PBS. HEK-293T cells were cultured on cover glass and transfected with pAAV-hSyn-RCaMP alone or combined with pAAV-hSyn-Nb-FT-2-TRPV1^Ca2+^ or pCMV-TRPV1^Ca2+^. Cells were then placed in glass bottom dishes with calcium imaging buffer at 29-31°C for imaging. Imaging was performed using a Deltavision personal DV imaging system (Applied Precision, Inc., Issawaq, WA) equipped with a custom-made ceramic lens and softWoRx imaging station. All other calcium imaging experiments were performed on an inverted Zeiss Axio Observer Z1 microscope (Carl Zeiss, Oberkochen GER). Cells were imaged before and during RF treatment, before and during application of a permanent magnet, before and after treatment with capsaicin (1 μM, Sigma). Imaging was performed on at least three occasions for each condition. Image analysis was performed using Image J. Briefly regions of interests (ROI) were selected for background or cells and mean intensity was calculated for every ROI in each image. Calcium responses were quantified as fluorescence intensity normalized to baseline fluorescence.

#### Immunocytochemistry of Cultured Cells

Immunocytochemistry (ICC) and immunohistochemistry (IHC) were used to confirm cell surface expression of TRPV1. Live cells were incubated with rabbit anti-TRPV1 (extracellular) polyclonal antibody (Thermo Fisher Scientific cat#PA5-77361) (1:50) in culture medium for 10 min at 37 °C followed by 5 washes with RT culture medium. Cells were then incubated with goat anti-rabbit Alexa Fluor 568 or 633 (1:1000,) for 10 min at 32 °C followed by a further 5 washes with RT culture medium. Cells were then fixed in 3.7% paraformaldehyde in Hank’s balanced salt solution (HBSS) for 30 min at room temperature and washed 5 times in HBSS before mounting using Fluoromount with DAPI (Southern Biotech, Birmingham, AL). Images were acquired using a Zeiss LSM 880 inverted confocal microscope (Carl Zeiss, Oberkochen GER).

### Electrophysiology recordings of N2A cells expressing pAAV-hSyn-Nb-FT-2-TRPV1^Ca2+^ construct

Neuro2A cells were cultured on uncoated petri dishes and were transfected with pAAV-hSyn-Nb-FT-2-TRPV1^Ca2+^ (as above). Electrophysiological recordings were performed 24-48 hours after transfection in a bath solution consisting of 10mM Hepes p7.4, 140 mM NaCl, 5 mM EGTA and were imaged using a Nikon eclipse DIC microscope at x20 magnification. Pipettes of borosilicate glass (Sutter Instruments; BF150-86-10) were pulled to ∼3-5 MW resistance with a micropipette puller (Sutter Instruments; P-97) and polished with a microforge (Narishige; MF-83). The pipette was filled with identical bath solution. Recordings were obtained with an Axopatch 200B amplifier (Molecular Devices), filtered at 1 kHz and digitized at 10 kHz (Digidata 1440; Molecular devices). Recordings were made in the outside-out excised patch configuration after gigaseals were obtained and currents were recorded by voltage ramp protocols from -100 to +100mV with and without 10 mM capsaicin addition by perfusion (ALA Scientific VM8 manifold perfusion system).

### Generation of AAV Constructs

pAAV-JET-DIO-MCS/pLP362 is the backbone for the AAV vectors used to make **pAAV-JET-DIO-mCherry, pAAV-JET-DIO-Nb-FT-TRPV1^Ca2+^, pAAV-JET-DIO-Nb-Ft-TRPV1^Cl-^ and pAAV-JET-DIO-TRPV1^Ca2+^** and was made by replacing the BsrGI/NheI region of pAAV-hSyn-DIO-hM3D(Gq)-mCherry (Addgene Cat No. 44361) with a multiple cloning site (TGTACAACTAGTACCGGTTCGCGAGCATGCCCTAGGGCTAGC) and inserting a 242-bp PCR product containing the JET promoter(*56*) amplified using primers 5’-CTGCGGCCGCACGCGTGTACCATTGACGAATTCGGGCG and 5’-ACTCTAGAGGATCCGGTACCTGTCAAGTGACGATCACAGGG into the MluI/KpnI sites. To make **pAAV-JET-DIO-mCherry/**pLP388, the mCherry open reading frame was amplified using primers 5’-ATTACCGGTGTTAACATGGTGAGCAAGGGCGAGGA and 5’-TAAACCGGTCTTAAGTTACTTGTACAGCTCGTCCATG and cloned into AgeI-digested pAAV-JET-DIO-MCS with the mCherry ORF in the reverse orientation to the JET promoter.

**pAAV-JET-DIO-Nb-FT-TRPV1^Ca2+^/**pLP363 was made by ligating a 2943bp SpeI/EcoRV fragment containing NbFT-TRPV1^Ca2+^ (*^34^*) to NruI/AvrII digested pAAV-JET-DIO-MCS. The 865-bp HindIII/Tth111I fragment of pAAV-JET-DIO-Nb-FT-TRPV1^Ca2+^ was replaced with a 445-bp HindIII-Tth111I fragment of synthesized DNA (Azenta Life Sciences, NJ) lacking the Nb-Ft sequence to make **pAAV-JET-DIO-TRPV1^Ca2+^**/pLP452.

To make **pAAV-JET-DIO-Nb-Ft-TRPV1^Cl-^/**pLP375, primers 5’-TCCATGGTGTTCTCCCTGGCAATGGGCTGGACCAACATGCTCT and 5’-AGACTAGTGTTATTTCTCCCCTGGGACCA were used to remove one NcoI site and amplify an 876-bp NcoI fragment of TRPV1^mutant^ encoding the I679K mutation^10^ which was cloned into NcoI-digested pAAV-JET-DIO-NbFT-TRPV1^Ca2+^.

pAAV-CMV-Cre-GFP was obtained from Addgene Cat No. 68544.

### Cell Culture and Adeno-Associated Virus (AAV) Preparation

Human embryonic kidney (HEK293, ATCC #CRL-1573) cells were cultured in DMEM (Gibco), supplemented with 10% (vol/vol) FBS (Sigma-Aldrich) and 1% (vol/vol) penicillin-streptomycin (Gibco), at 37 °C in 95% humidified air and 5% CO2. Vector stocks were prepared by packaging the plasmids into rgAAV2 or mixed serotype AAV1/2 particles using a calcium phosphate transfection system as described previously(*57*). Cells were harvested and lysed at 72 h after transfection. The vectors were purified using iodixanol gradient and dialyzed against PBS with 2mM MgCl2. AAV titers were determined by qPCR using Syber Green chemistry and with primers to the WPRE fragment of the AAV backbone.

### Quantitative Real Time PCR (qPCR)

To titer our purified viral vectors, AAV were processed as previously described(*57*) and viral genomes quantified via qPCR using Fast SYBR green master mix (Applied Biosystems Cat. No 4385612) on the AB 7500 FAST Real Time PCR platform (Applied Biosystems) and primers to the WPRE element: WPRE-Fw: 5’-GGCTGTTGGGCACTGACAAT-3’; WPRE-Rev: 5’-CTTCTGCTACGTCCCTTCGG-3’. The relative number of full viral particles were calculated using the standard curve method by normalizing to known standard samples.

### Animals

Male mice were housed two to five per cage and kept at 22°C on a reverse 11 am-light/11 pm-dark cycle, with standard mouse chow and water provided ad libitum throughout the duration of the study. All animal procedures were approved by the Institutional Animal Care and Use Committee of Weill Cornell Medicine and were in accordance with National Institutes of Health guidelines. The following mice were used in these studies: i) wt C57BL/6J mice, obtained from Jackson laboratory (Jax.org; strain #:000664); ii) 129S-Pitx2tm4(cre)Jfm/Mmucd (Pitx2-Cre; MMRRC stock #:000126-UCD) and *B6.FVB(Cg)-Tg (Adora2a-cre) KG139Gsat/Mmucd* (A2a-Cre; MMRRC stock #:036158-UCD).

### Stereotactic Surgery

All stereotactic surgical procedures were performed on 8–12-week-old mice, weighing 20g to 25g, under a mixture of ketamine/xylazine anesthesia. Ketamine (Butler Animal Health Supply) and xylazine (Lloyd Laboratories) were administered at concentrations of 110 and 10 mg/kg body weight i.p., respectively. After the induction of anesthesia, the animals were placed into a stereotactic frame (David Kopf Instruments). All infusions were performed using a 10-μL stereotactic syringe attached to a micro-infusion pump (World Precision Instruments) at a rate of 0.1-0.4 μL/min. To prevent reflux, after each infusion, the injection needle was left in place for 5 min, withdrawn a short distance (0.3–0.5 mm), and then left in the new position for an additional 2 min before removal. With a 33 G needle To generate 6OHDA lesioned mice, animals were injected with a total volume of 0.6 μL of 6OHDA hydrobromide (Sigma-Aldrich) in PBS with 0.1% ascorbate unilaterally into the medial forebrain bundle (MFB) at a concentration of 2.5 mg/mL and an infusion rate of 0.1 μL/min with a 10-μL Hamilton syringe with a 30 G needle. The coordinates for the injection were AP −1.1 mm, ML -1.1 mm, DV −5.0 mm relative to bregma and the dural surface. Before lesion surgery, the norepinephrine reuptake inhibitor desipramine (25 mg/kg, i.p.) was administered via i.p. injection at least 30 min before 6OHDA infusion, to protect neostriatal and cerebellar noradrenergic neurons from the toxin-induced damage. Mice were allowed 4-6-week recovery before being subjected to the apomorphine-induced a behavioral test to estimate the extent of dopamine depletion in the substantia nigra (see following section).

For experiments in Pitx2-cre mice, AAV1/2 vectors were injected into the subthalamic nucleus (STN) unilateral to the lesioned side (AP -2.0 mm, ML -1.7 mm, DV −4.7 mm from bregma) of 6OHDA mice. For experiments in A2a-cre mice, AAV1/2 vectors were injected bilaterally into the dorsal striatum (AP +0.5 mm, ML ±2.3 mm, DV −3.5 mm from bregma). For experiments in wt mice, AAV1/2 vectors were injected bilaterally into the dorsal striatum (AP +0.5 mm, ML ±2.3 mm, DV −3.5 mm from bregma) and AAV2retro into the globus pallidus (AP +0.5 mm, ML ±2.3 mm, DV −3.5 mm from bregma). All craniotomies to inject AAV vectors were performed at a rate of 0.4 μL /min using a 10-μL WPI syringe with a 33 G needle. A total of 2x10^11^ genomic particles/μL (2 μL in PBS) of vectors were injected.

The mice were allowed a 6–8-week recovery before being subjected to the behavioral tests during which maximal transgene expression from AAV vectors is achieved. At the conclusion of the experiments, injection site accuracy within the targeted brain region was determined by immunohistochemistry for the specific transgene expressed by the AAV vectors, and mice with mistargeted injections were excluded from analysis before their data were unblinded.

### Sources of Magnetic field

#### Direct magnetic field (DMF)

The magnetic field for *in vivo* studies was generated by the superconducting electromagnetic MRI field from a Siemens 3.0 Tesla PRISMA MRI Scanner (Siemens Healthcare). The magnetic field strength was measured and mapped using a gaussmeter (F.W. BELL MODEL 5080). High DMF was defined as a region with a strength of 0.5-1.3 T, and Low DMF as 0.1-0.27 T.

#### Transcranial magnetic stimulation (TMS)

A butterfly C-B60 coil (165 x 85 x 19mm/inner diameter 35mm/outer diameter 75mm) was used to deliver magnetic field during *in vivo* studies. Biphasic stimulation pulses were delivered by a waveform generator (MagPro100x with MagOption) with the following settings: 20% machine capacity (∼600 mT), twin/dual mode 50 Hz stimulation, 500μs rise time, 1ms inter pulse interval, 10 pulses in train, number of trains 20, and inter train interval 1 s for 20 s. This set up was determined based on trials where small percent increments in machine capacity were performed until a freezing phenotype was observed or until an electric shock-induced response was recorded. In those trials, 25% of TMS capacity caused electric shock-induced responses such as twitching (**TMS method supplementary videos**).

### Behavioral Assessments

All behavioral tests were run during the dark phase of the mouse daily cycle. The experiments were performed by an examiner blinded to treatment groups.

#### Apomorphine induced rotations

To test for 6OHDA lesion efficiency in Pitx2-cre mice, rotational behaviors were performed. In brief, mice were placed in a body harness connected to a transducer/swivel in a bowl-shaped testing arena (rotometer). Full 360° clockwise and counter-clockwise rotations were measured over a 45-min period after drug injection for apomorphine-induced rotations. Apomorphine (Sigma-Aldrich) was dissolved in saline solution, 0.9% NaCl, and 0.1% ascorbate, and administered at a dose of 0.25 mg/Kg via s.c. injection. Mice we randomized based on the net number of contralateral rotations before receiving STN injections of the inhibitory magnetogenetic construct.

To test the effect of our AAV.Nb-Ft-TRPV1^Cl-^ vectors on inhibiting STN neuronal activity, rotational behaviors were performed inside a 3T MRI room in a sound insulated, ventilated Styrofoam chamber that contained a plastic arena of 26.5 x 26.5 x 26.5 cm. Video recordings of apomorphine induced rotations were obtained beginning 10 minutes after 0.25 mg/Kg via s.c. injections for 5 minutes outside the magnetic field followed by 5 minutes inside the magnetic field [1.3T – 500mT], and another 5 minutes outside the magnetic field in the start position using a magnetic field compatible camera (MRC HiSpeed camera, https://www.mrc-systems.de/en/products/mr-compatible-cameras#12m-camera). Upon the conclusion of behavioral studies, nigral lesion efficiency by 6OHDA was determined by immunohistochemistry for the tyrosine hydroxylase (TH) protein as a marker for dopaminergic neurons in the substantia nigra. Mice with mistargeted injections were excluded from analysis before their data were unblinded.

#### Open-Field Arena

Locomotor behavior in an open-field arena was performed inside a 3T MRI room using the same chambers and video recording system as described above for apomorphine induced rotations. For 2 days prior to testing, each mouse was habituated in the open field box for 10 minutes to reduce any possible confounding effect of a new environment. On the day of testing, each mouse was placed in the center of the open field, and spontaneous horizontal and vertical behaviors were recorded during 5-min sessions of pre-DMF, during DMF, and post-DMF.

#### Noldus EthoVision

EthoVision XT14 was used to analyze offline rotational behaviors and open field videos. The automatic animal detection with deep learning settings was used to track center to-nose and center to-tail points in defined arenas. Video recordings were used to analyze the following locomotion and rotational parameters: (1) distance travelled, (2) activity percentage (%), (3) rotation >90 degrees clockwise, and counterclockwise. Tracking data and heatmaps were graphed with EthoVision analysis tools.

#### TRACKER

TRACKER video analysis and modeling tool (TRACKER 6 free software, https://physlets.org/tracker/) was used to track precise motor responses, including freezing and ambulation time. Each second of the recordings was divided into 5 frames to track nose, center, fore and hind paws, and tail every 2 frames resulting in 150 positions measured per minute. Spatial motion in the Y axis, referred as “change of location”, was also analyzed as a function of time.

### Immunohistochemistry of Brain Sections

On completion of all assessments, mice were deeply anesthetized with sodium pentobarbital (150 mg/kg, i.p.) and transcardially perfused with 4% paraformaldehyde (PFA). Brains were extracted and post-fixed overnight in 4% PFA, cryoprotected in 30% sucrose, and cut into 30-μm sections using a vibratome (Leica Microsystems). Free-floating sections were treated with various antibodies to visualize proteins of interest using immunofluorescence labeling, including tyrosine hydroxylase (TH) to quantify the extent of nigral neurodegeneration, and mCherry or HA to estimate the transduction efficiency from our AAV vectors. The following primary antibodies were used: rabbit anti-RFP to detect mCherry (Rockland Cat. No 600-401-379; dil. 1:1000), sheep anti-TH (Millipore Cat. No AB1542; dil. 1:1000), rabbit anti-HA (Sigma-Aldrich Cat. No H6908, dil.1:500), and rat anti-cfos (Synaptic systems Cat. No 226-017, dil. 1:1000). Alexa Fluor-conjugated fluorescent secondary antibodies (Molecular probes) were obtained from Life Technologies and used at a 1:1000 dilution. For *cfos* analyses, mice were exposed to the DMF (MRI magnet 500mT -1.3T) for 20 minutes prior being transcardially perfused as described above. All mice that were included in the *cfos* data were perfused within one hour following DMF exposure.

### RNAScope in situ hybridization

Thirty μm free floating brain sections were mounted in PBS onto charged Superfrost slides (Fisher Scientific) and were allowed to air dry for ∼ 2h. The experiments were performed as per manufacturer’s instructions (ACDbio). The incubations at 40°C and 60°C took place in the HybEZ oven using the EZ-Batch system (ACDbio). Sections were first baked at 60°C for 1h and post-fixed in 4% PFA for 15 min at 4°C. After the dehydration steps, slides were dried at 60°C for 15 min, treated with hydrogen peroxidase (Cat No. 322335, ACDbio) at room temperature for 10 min, and dried again at 60°C for 15 min before performing the retrieval step. Sections were incubated in target retrieval buffer (Cat No. 322000, ACDbio) for 10 min at 100°C and washed two times with double deionized water (ddH20). After 3 min incubation in 100% Ethanol and drying at 60°C for 10 min, sections were then incubated in Protease III (Cat No. 322337, ACDbio) at 40°C for 30min, washed five times with ddH20, and incubated for 2h at 40°C with probes including Mm-Drd2 – C1, Mm-Drd1 – C3, mCherry-C4. A slide for either positive and negative control probes were also included (Cat No. 321811 and 321831 respectively, ACDbio). Two washes occurred in between the following 40°C incubations with wash buffer (Cat No. 310091). Sections were incubated for 30min with AMP1, 30 min with AMP2, and 15min with AMP3 (kit Cat No.323110). Next, sections were incubated in HRP C1 for 15min and Opal 520 (1:1500; Cat No. OP-001001) in TSA buffer (Cat No. 322809). Sections were next incubated in an HRP blocker for 15 min. Following sections were incubated with HRP C2 in combination with Opal 650 (1:1500; Cat No. OP-001003), and HRP C4 in combination with Opal 570 (1:1500; Cat No. OP-001005). Slides were cover slipped with Prolong Gold containing DAPI (ACDbio).

### PET/CT animal imaging studies

All animal imaging experiments were performed in the Citigroup Biomedical Imaging Center (CBIC, Weill Cornell Medicine). For baseline experiments, mice were fasted for 12 hours prior imaging followed by. Intraperitoneal (i.p.) injection of ∼500μCi ^18^F-fluorodeoxyglucose (FDG), and residual dose was measured. 1 hour after injection, mice were anesthetized with isoflurane and scanned for 1 hour in a micro-PET/CT scanner (Siemens Inveon). For DMF experiments, mice were exposed to a 3T MRI (DMF=500 mT – 1.3 T) for a total of 25 minutes after ^18^F-FDG i.p., administration, and scanned following the same protocol as for baseline experiments. Following imaging, scans were histogrammed and reconstructed using Siemens Inveon Acquisition Workplace (Siemens Medical Solutions, Knoxville, TN, USA) and analyzed in DICOM format.

MRICron and MATLAB were used to do a PET-CT co-registration, and PMOD Fusion tool (version 3.6, PMOD Technologies Ltd., Zurich, Switzerland) was used for rigid transformations (manually co-registered via rigid transformations [translations and rotations]), matching with MRI mouse template, normalization with Mirrione mouse brain atlas, and regions of interest (ROI) analysis. All analyses were conducted in a blinded fashion and were adjusted for mouse weight, ^18^F-FDG final dose (^18^F-FDG_initial dose_-^18^F-FDG_residual dose_), as well as relative to regions that do not express dopamine 2-type receptors such as cerebellum. Right and left dorsal striatum were averaged for each animal and MBq/ml or Percent injected dose per gram (%ID/g) were calculated to plot data.

### In vivo fiber photometry

Fiber photometry was performed to measure *in vivo* calcium dependent activity dynamics during DMF/TMS stimulation. Cre-dependent AAVs expressing GCaMP6s (AAV9-Syn-DIO-GCaMP6s, Addgene Cat. No 100845-AAV9) and Nb-Ft-TRPV1^Ca2+^-HA/mCherry were injected bilaterally into the dorsal striatum of A2a-Cre transgenic mice to restrict expression of GCaMP6s and NbFt- TRPV1^Ca2+^-HA or mCherry to D2 iSPNs. Two weeks following AAV injection, a 400-μm-diameter optical fiber (Doric, MFC_400/430-0.48_5.5mm_ZF2.5(G)_FLT, Cat. No B280-4418-5) was implanted in the dorsal striatum 0.5 mm above the AAV injection site (AP: +2.00mm, ML: −0.25 mm, DV: −2.15mm) and was secured with Metabond. Prior to behavioral testing, mice were habituated for 2 days to a patch cord attached to the implanted optical fiber for 3 min sessions. During behavioral testing, fiber photometry recordings were taken for 2 min before, during, and post magnetic stimulation. Behaviors were performed and recorded using the same settings as described above for open-field arena locomotor quantification. Following behavioral testing, fluorescence microscopy was utilized to confirm GCaMP6s expression and optical fiber placement. To excite GCaMP6s, light from a 470nm LED (Thorlabs, M470F3) modulated at a frequency of 521Hz was passed through a filter (Semrock, FF02-472/30), reflected by a dichroic (Semrock, FF495-Di03) and coupled to a 0.48 NA, 400µm core optical fiber patch cord (Doric). Emitted fluorescence traveled back through the patch cord, passed through the dichroic, a filter (Semrock, FF01-535/50), and was focused onto a photodetector (Newport, Model 2151). The modulated signal passed from the photodetector to a RP2.1 real-time processor (Tucker Davis Technologies) where it was demodulated, and low pass filtered using a corner frequency of 15Hz. TTL pulses denoting the start of behavioral trials were passed to the processor in real time for alignment of calcium recordings to behavioral measures.

### In vitro chloride imaging

Neuro 2A cells (ATCC ccl-131, mycoplasma testing and STR profiling performed by ATCC) were grown in Eagles’ minimum essential medium with 10% fetal bovine serum (Gibco) at 37°C and 5% CO_2_.

For immunocytochemistry studies, cells were cultured on 12-mm cover glass (Fisher Scientific, Pittsburgh, PA) coated with fibronectin (Sigma). Cells were transfected with constructs expressing Nb-Ft-TRPV1^Cl-^ or TRPV1^Ca2+^ 24 h after plating. Cells were stained 48-72hrs after transfection. For imaging studies using the chloride-dependent reporter MQAE, Neuro2A cells were grown on 12-mm cover glass and transfected with plasmids expressing mCherry alone, co-transfected with NbFt-TRPV1^Cl-^ and mCherry or co-transfected with TRPV1^Ca2+^ and mCherry. Holotransferrin (2 mg/ml, Sigma) was added to cells 24 hours after transfection. Cells were placed at 32°C 24 hours before imaging. Cells were washed 3 times with Krebs-HEPES buffer (NaCl 110 mM, KCl 5.5 mM, CaCl_2_ 2.5 mM, MgCl_2_ 1.25 mM, HEPES 5mM and glucose 10mM) then incubated with MQAE 5mM (Invitrogen) in Krebs-HEPES buffer for 1 hour at 37°C. Cells were then washed in Krebs-HEPES buffer three times and placed in glass-bottom dishes with Krebs-HEPES buffer for 15 minutes before imaging. Imaging was performed using an inverted fluorescent microscope Zeiss Axio Observer Z1 microscope (Carl Zeiss, Oberkochen GER). Cells were imaged every 6 seconds for a total of 3 mins with imaging before and during low AMF treatment with a neodymium magnet (K and J magnetics) to produce a peak field 230-250mT. Image analysis was performed using FIJI. Briefly, regions of interest (ROI) were selected based on mCherry expression and mean intensity calculated for each ROI in each image after background subtraction. Chloride responses were quantified as change in fluorescence intensity normalized to baseline fluorescence (ΔF/F0). Imaging was performed on at least three occasions for each condition and chloride responses were measured in 142 cells (Nb-Ft-TRPV1^Cl-^), 76 cells (TRPV1^Ca2+^) and 181 cells (mCherry).

Immunocytochemistry (ICC) was used to confirm cell surface expression of TRPV1. Live cells were incubated with rabbit anti-TRPV1 (extracellular) polyclonal antibody (Thermo Fisher Scientific cat#PA5-77361) (1:50) in culture medium for 10 min at 37 °C followed by 5 washes with RT culture medium. Cells were then incubated with goat anti-rabbit Alexa Fluor 633 (1:1000) for 10 min at 32 °C followed by a further 5 washes with RT culture medium. Cells were then fixed in 4% paraformaldehyde in Hank’s balanced salt solution (HBSS) for 30 min at room temperature and washed 5 times in HBSS before mounting using Fluoromount with DAPI (Southern Biotech, Birmingham, AL).

### Imaging acquisition and analysis

Images were taken by an epi-fluorescent microscope (Olympus BX61 microscope fluorescent microscope fitted with an Olympus DP71 digital camera) or a confocal microscope (Zeiss LSM 900) and analyzed with Image J software (version 1.52p, NIH, US). For quantification of *c-fos*+ cells, coronal sections were sampled at intervals of 120-160μm for immunostaining. 3 dorsal striatal or nigral sections per mouse were identified using the mouse brain atlas of Paxinos (2007) and scanned bilaterally using a 20x objective. For each selected section, three randomly chosen regions of interest (ROIs) with fixed areas were selected for quantification. A Macro Plugin was applied to each section obtained, with a Gaussian blur filter threshold set to 30 and size filter to 20 to remove small objects and low signal cells. *c-fos*+ neurons were determined only when clear co-localization with DAPI staining was observed. Automatized detection using Image J software in 4-5 mice per condition (controls vs treated).

### Statistics

To make valid comparisons among experimental groups, mice were randomized based on baseline motor behavior to achieve balanced, unbiased groups. Pitx2-cre mice were randomized based upon contralateral rotational counts following systemic apomorphine injection (0.25 mg/Kg), measured at 4 weeks after 6OHDA lesion surgery. The sample size for our experiments was determined by power analysis(*58*), published work in the field, and previous studies performed by the PIs. Exact sample sizes for each experiment are indicated in figure legends. The power was set at 95% probability, effect size to 0.3-0.5 depending on the experimental outcome, and significance at p<0.05.

Statistical analyses were conducted using GraphPad Prism 7.0. The statistical test used in each experiment depended on the type of data collected (reported in figure legends). We tested for normality using the Kolmogorov-Smirnov test and Q-Q plots. Two-way ANOVA or two-tailed t test for statistical comparisons were performed as appropriate.

For fiber photometry experiments, the change in fluorescent intensity (ΔF/F) percentage was calculated by subtracting the linear fit of the isosbestic signal to the raw GCaMP signal from the raw GCaMP signal, median for the entire recording session was then subtracted from each GCaMP event value and divided by the median; that is, (465_raw_ – 405_fitted_) = 465_net_; (465_net_ – 465_median_)/(465_median_) x 100. Events were computed comparing means in pre-DMF/DMF/post-DMF or baseline/TMS conditions.

Pearson’s correlation coefficients were used to quantify associations between different readouts (e.g., c-fos+ immunoreactivity vs freezing of gait). When comparing two sets of normally distributed data, unpaired two-tailed t-tests were utilized.

## Acknowledgments

We acknowledge Henning U. Voss and Immanuel Elbau for assistance with the video-recording systems during MRI and TMS, William Tower for assistance with the EthoVision software, Neranjan de Silva for assistance breeding mice, Violetta Manusajyan for technical support during MRI recordings and Muc Chieu Du and Kamaar Lawson for assistance with use of the Citigroup Biomedical Imaging Center.

## Author contributions

Conceptualization: JMF, MGK

Methodology: SRU, LEP, XY, RM, LK, JMF, MGK

Investigation: SRU, LEP, XY, LK, GH, HM, GV, PW, AMA, SN, JPD, EKF, LG, RZ, SAS, MGK

Visualization: JMF, MGK. Funding acquisition: MGK, JMF

Supervision: SAS, JMF, MGK

Writing – original draft: SRU, MGK

Writing – review & editing: SRU, SAS, JMF, MGK

## Data and materials availability

All data are available in the main text or in the supplementary materials. For additional information corresponding author should be contacted, Michael G. Kaplitt

## Supplementary Figure Legends

**Supplementary Fig. 1. Generation of Nanobodies (Nb) to human and mouse ferritin. A.** Schema of overall method for llama immunization, generation of phage display library, panning for selection of highly reactive nanobody clones to human spleen ferritin and mouse liver ferritin, ELISA and numbers of unique nanobodies reactive to human, mouse or both human and mouse ferritin. Absorbance ratio of nanobody clones for human spleen ferritin (**B, C**) and mouse liver ferritin (**D**). **E**. Classification of nanobody clones based on their similarity on their complementary region 3 (CD3). **F**. Example of ELISA selection for nanobodies from distinct groups for highly reactive clones. Clones were screened after the 1^st^ and 2^nd^ rounds of panning against human ferritin and after the 2^nd^ round of panning against mouse ferritin. Clones exhibiting absorbance ratio greater than 2.5 (dashed red line) were selected for further characterization. **G**. Alignment of amino acid sequence of nanobodies belonging to CDR3 groups 6 and 26 with the highest reactivity to human and mouse ferritin. CDR1, CDR2, and CDR3 are colored in blue. The nanobodies’ CDR3 group and names are on the right. **H**. Schema of method for quantification of nanobody binding to human spleen ferritin. HA-tagged nanobodies from CDR3 groups 6 and 26 were purified from transformed WK6 E. coli by metal-ion affinity chromatography.

**Supplementary Fig. 2. Screen of Nb clones to mouse ferritin. A.** Schema of method to characterize Nb-Ft binding to mouse ferritin. Mammalian expression plasmids with Nb-Ft sequences (Nb-Ft-2, -9, -10, -14, -17) or Nb-GFP fused to HA-tagged mCherry were co-transfected into HEK-293T cells along with a plasmid expressing GFP-tagged mouse ferritin (GFP-mFerritin). After 48 hours, cells were lysed and HA-tagged nanobodies along with their bound proteins were immunoprecipitated using anti-HA conjugated agarose beads. Eluted proteins were examined by SDS-PAGE and mass spectrometry. **B.** Quantification of nanobody-bound proteins by mass spectrometry normalized for protein signal (log2). The protein signals for mouse ferritin heavy chain (transfected as GFP-mFerritin), human ferritin heavy chain (FTH1_human, endogenous to HEK-293T cells), and N-terminal GFP moiety of GFP-mFerritin are shown along with a control peptide, human tubulin (TBB5_human). The overall protein signal shown in the box plot does not differ between test conditions. **C.** Schema of method to validate ability of Nb-ferritin clones to transduce magnetic field into cell activation. Constructs with Nb-Ft sequences (Nb-Ft-2, -9, -10, -14, -17) fused to the N-terminal of TRPV1^Ca2+^ were generated and transfected into HEK-293T cells along with a plasmid expressing GFP-tagged mouse ferritin (GFP-mFerritin) and a calcium-dependent secreted alkaline phosphatase (SEAP). Cells transfected Nb-GFP-TRPV1^Ca2+^/GFP-mFerritin and the reporter were used as a positive control. **D.** Schema of isothermal titration calorimetry assay for binding of anti-ferritin nanobody-2 (Nb-Ft2) to human spleen ferritin. Human spleen ferritin (10 μM) was titrated against 75 μM Nb-Ft2 (75 μM, single 0.4 μL injection, then 19 injections of 2 μL each, 150-sec intervals, 750 rpm stirring) with a reference power of 10 μcal/sec. **E.** Binding affinity and interaction between Nb-Ft2 and human spleen ferritin by isothermal titration calorimetry at 25°C (K_d_ = 0.54 ± 0.2 μM).

**Supplementary Fig. 3. Functional validation of Nb-Ferritin clones *in vitro.*** Cell surface expression of Nb-Ft-2-TRPV1^Ca2+^ by immunolabeling using antibodies directed to the extracellular portion of TRPV1 in transfected human HEK-293T (**A**) and murine Neuro2A cells (**B**) without permeabilization. Capsaicin-dependent currents in pulled patches from Neuro2A cells transfected with the hSyn-Nb-Ft-2-TRPV1^Ca2+^ construct (**C**) and untransfected cells (**D**). **E**. Changes in RCaMP fluorescence normalized to baseline fluorescence (ΔF/F_0_) with capsaicin treatment of HEK-293T cells expressing RCaMP alone (49 cells), TRPV1^Ca2+^ (65 cells) or Nb-Ft-2-TRPV1^Ca2+^ (73 cells). Error bars represent mean +/-SEM. **F**. Peak ΔF/F_0_ with capsaicin treatment of HEK-293T cells expressing RCaMP alone (49 cells), TRPV1^Ca2+^ (65 cells) or Nb-Ft-2-TRPV1^Ca2+^ (73 cells). Data were analyzed by ordinary one-way ANOVA with Tukey’s multiple comparison test, **** p < 0.0001. Error bars represent mean ± SD.

**Supplementary Fig. 4. Peak fluorescence in response to magnet treatment. A.** Peak ΔF/F_0_ of Neuro2A cells expressing Nb-Ft-2-TRPV1^Ca2+^ with (786 cells) or without (1659 cells) magnet treatment. Data were analyzed by Mann Whitney U test **** p < 0.0001. **B.** Peak ΔF/F_0_ with magnet treatment of HEK-293T cells expressing RCaMP alone (54 cells), TRPV1^Ca2+^ (107 cells) or Nb-Ft-2-TRPV1^Ca2+^ (101 cells). Data were analyzed by ordinary one-way ANOVA with Tukey’s multiple comparison test, **** p < 0.0001. Error bars represent mean ± SD. **Supplementary Fig. 5. Viral-mediated expression of Nb-Ft-TRVP1^Ca2+^ in striatal iSPNs elicit parkinsonian motor behavior. A.** RNA probe validation for D1 and D2 neurons**. B.** Example of Nb-Ft-TRVP1^Ca2+^ expressing mouse stained for HA (left), number of transduced neurons in STR between mCherry and Nb-Ft-TRPV1Ca2+ groups (upper right), and correlation of viral transduction with freezing of gait time during DMF (lower right).

**Supplementary Fig. 6. High DMF Titration in Nb-Ft-TRPV1^Ca2+^ expressing mice. A.** Schema for high direct magnetic field (DMF) titration. **B.** Magnetic field mapping in the “y” axis of the cage used for behavioral assessment. **C.** Individual activity heatmap example of motor activity during bilateral striatal High DMF titration in Nb-Ft-TRPV1^Ca2+^ expressing mouse. **D.** Freezing of gait in different magnetic field gradients in the High DMF titration ranges (red bars, n=5) in Nb-Ft-TRPV1^Ca2+^ expressing mice.

**Supplementary Fig. 7. Transcranial magnetic stimulation (TMS) treatment alters motor behavior in Nb-Ft-TRPV1^Ca2+^ expressing mice.** Activity percentage (%) during baseline (left) and TMS (right) treatment in the **A.** mCherry group (blue), and **B.** Nb-Ft-TRPV1^Ca2+^ group (red). **C.** Average distance during baseline and TMS treatment in mCherry (blue, n=4) and Nb-Ft-TRPV1^Ca2+^ (red, n=5) groups. Error bars show SEM. * represents p value < 0.05 with Student’s t test. Tracking data of change of position in the **D.** “y” axis, and **E.** “x” axis in mCherry (blue), and Nb-Ft-TRPV1^Ca2+^ group(red) during baseline and TMS treatment.

**Supplementary Fig. 8. Application of mutant Nb-Ft-TRPV1^Cl-^ membrane channel for subthalamic nucleus in mouse model of Parkinson disease**. **A.** Immunostaining for cell surface expression of TRPV1 in Neuro2A cells transfected with mCherry, TRPV1^Ca2+^ or Nb-Ft-TRPV1^Cl-^ (from left to right). Scale bar 20um. **B.** Representative immunostaining shows unilateral loss of TH+ neurons in the substantia nigra (SN) of lesioned PitX-2-Cre mice with 6-OHDA in the MFB. **C.** Randomization 6-OHDA lesioned mice based on apomorphine induced rotations. **D.** Selection of time points for behavioral testing during DMF application. Individual variation in the number of contralateral rotations for 5 minutes during pre-DMF, DMF, and post-DMF exposure in the **E.** mCherry group (n=10), and **F.** Nb-Ft-TRPV1^Cl-^ group (n=9).

