## Supplementary figures and images for "Bidirectional Regulation of Motor Circuits Using Magnetogenetic Gene Therapy"

### Supplementary figures

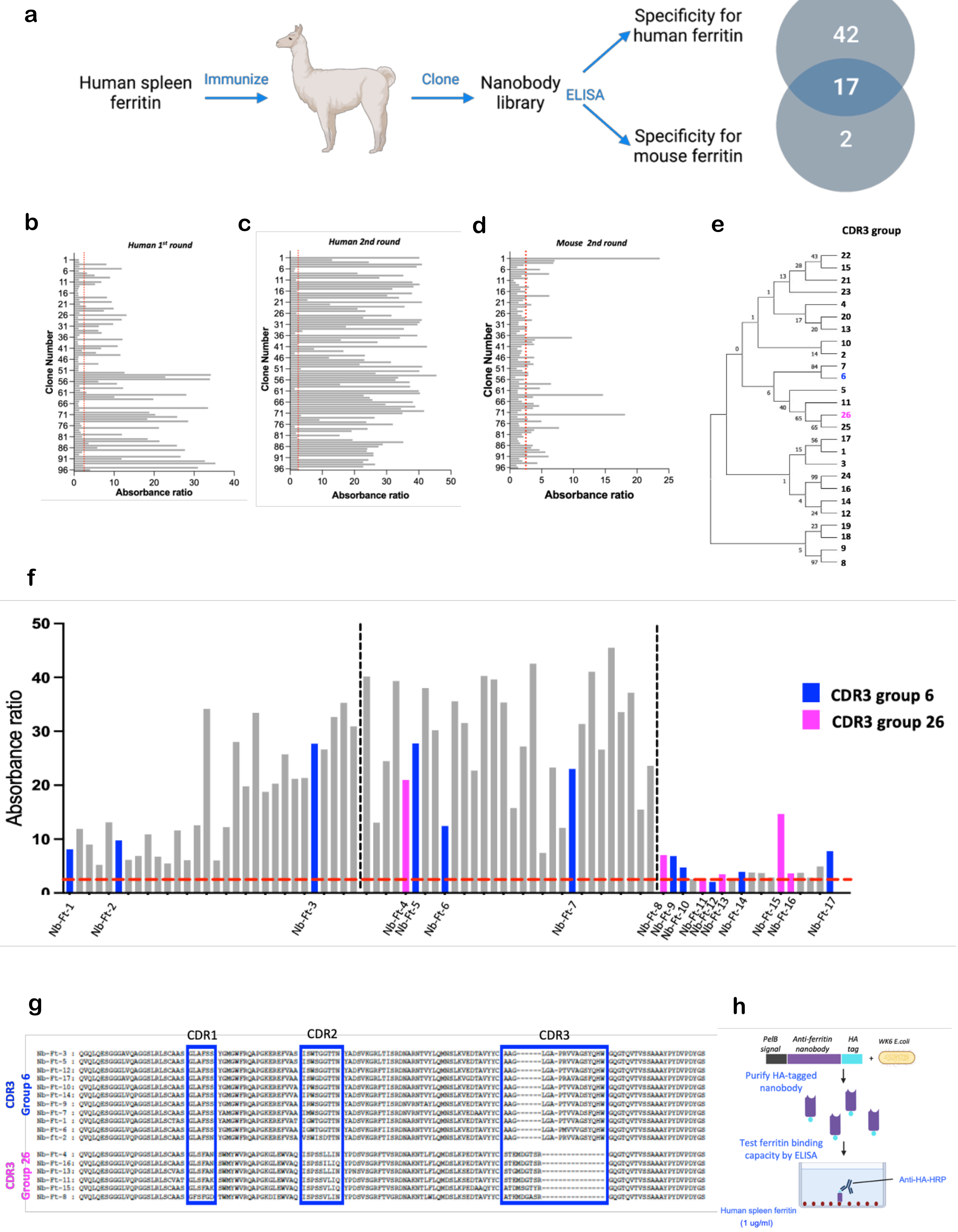

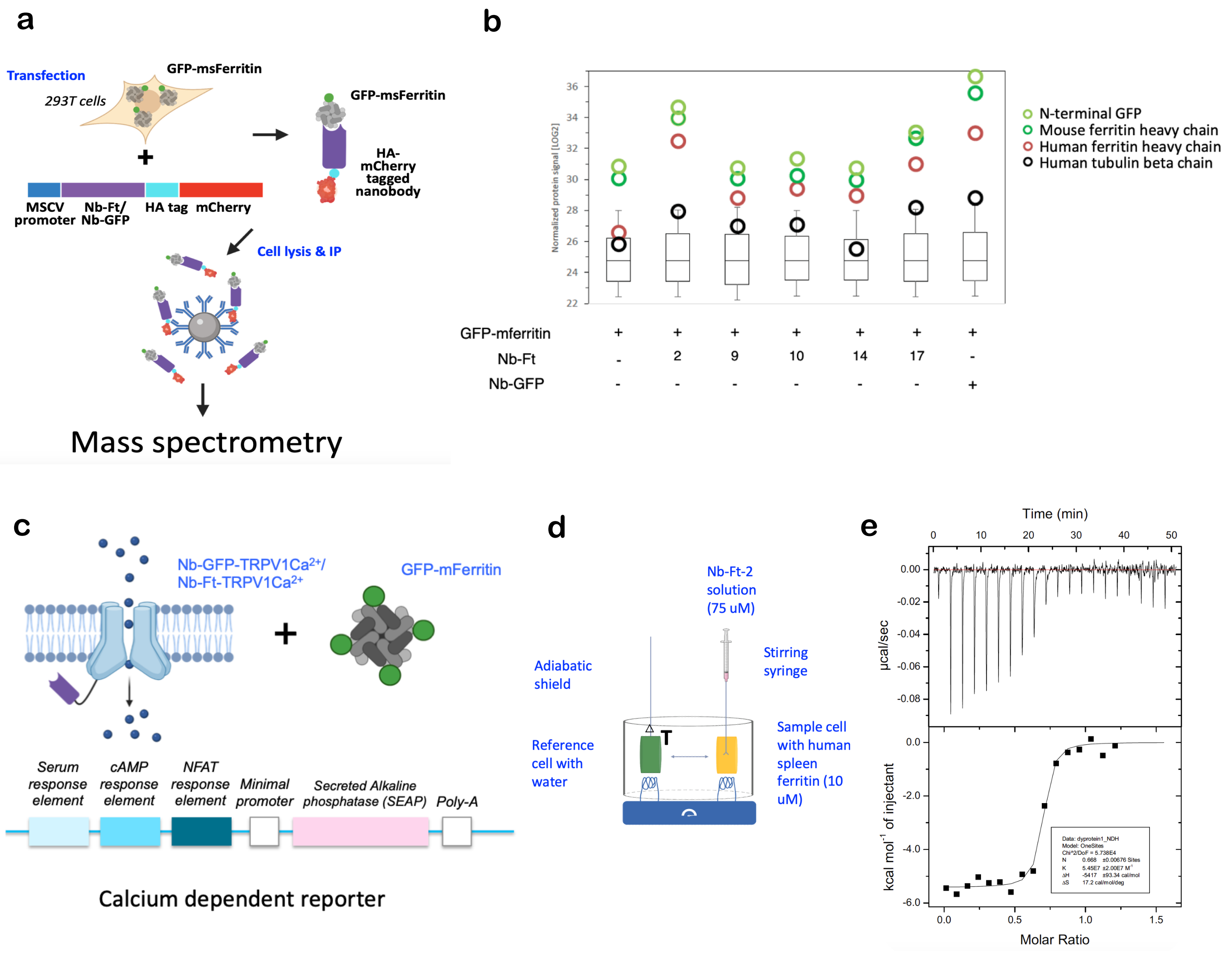

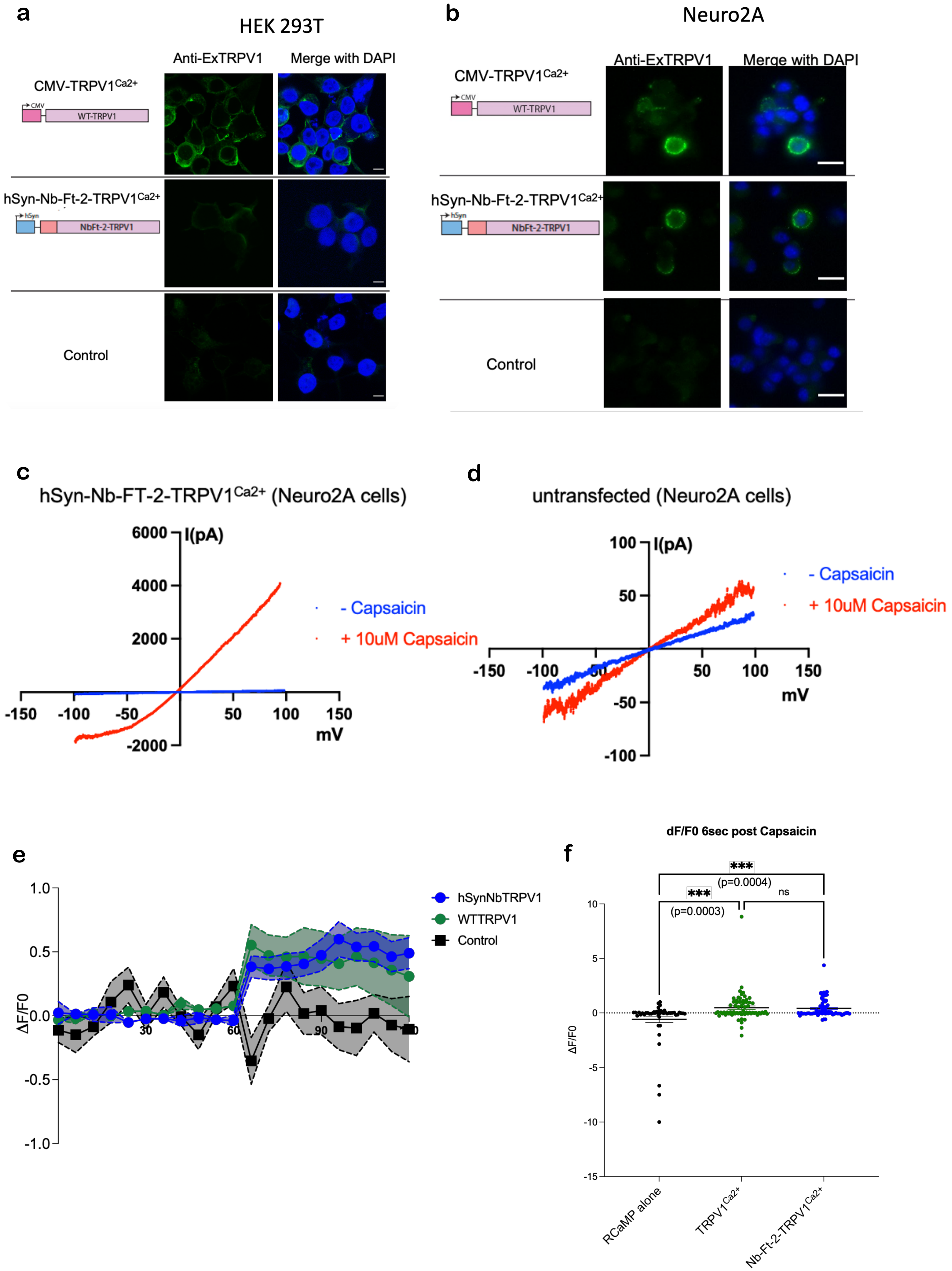

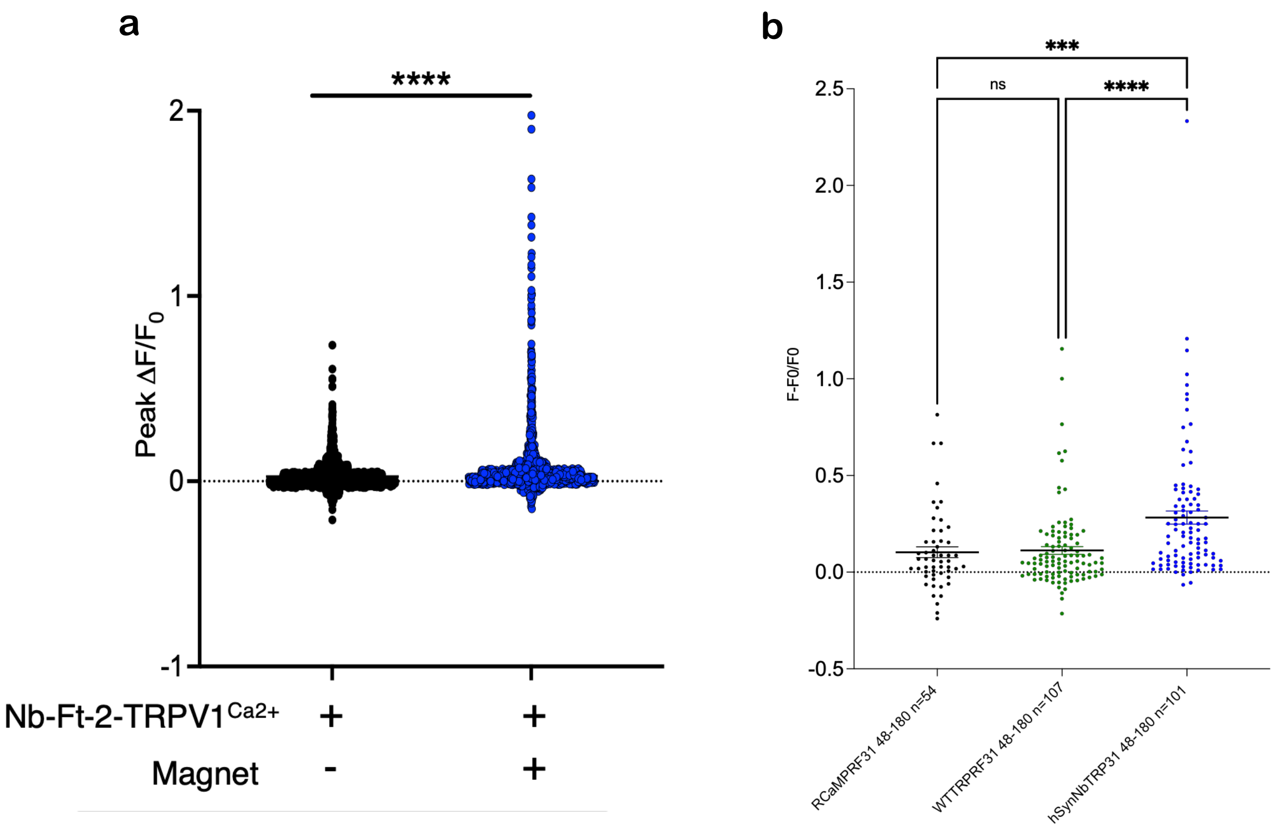

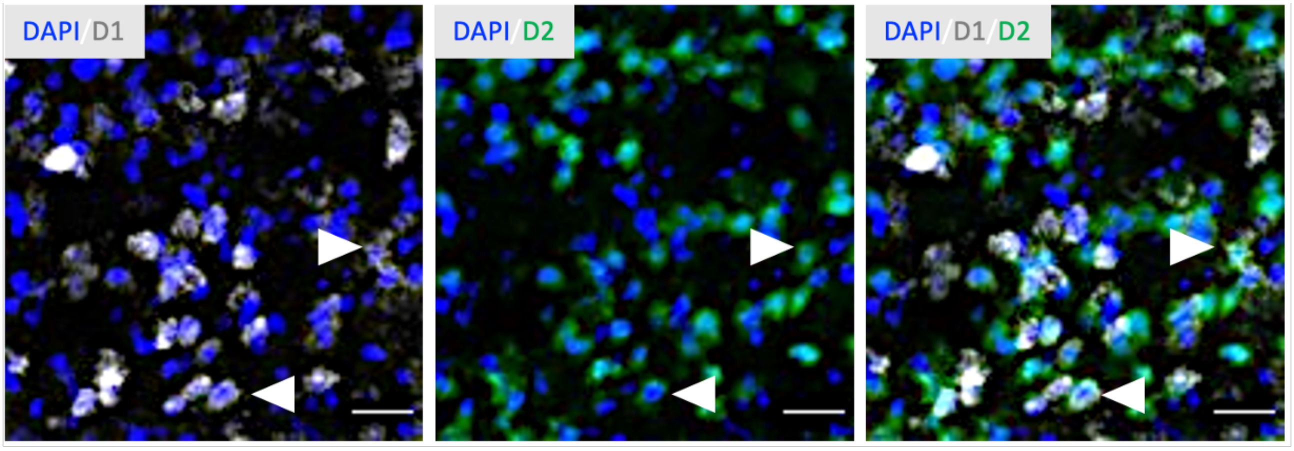

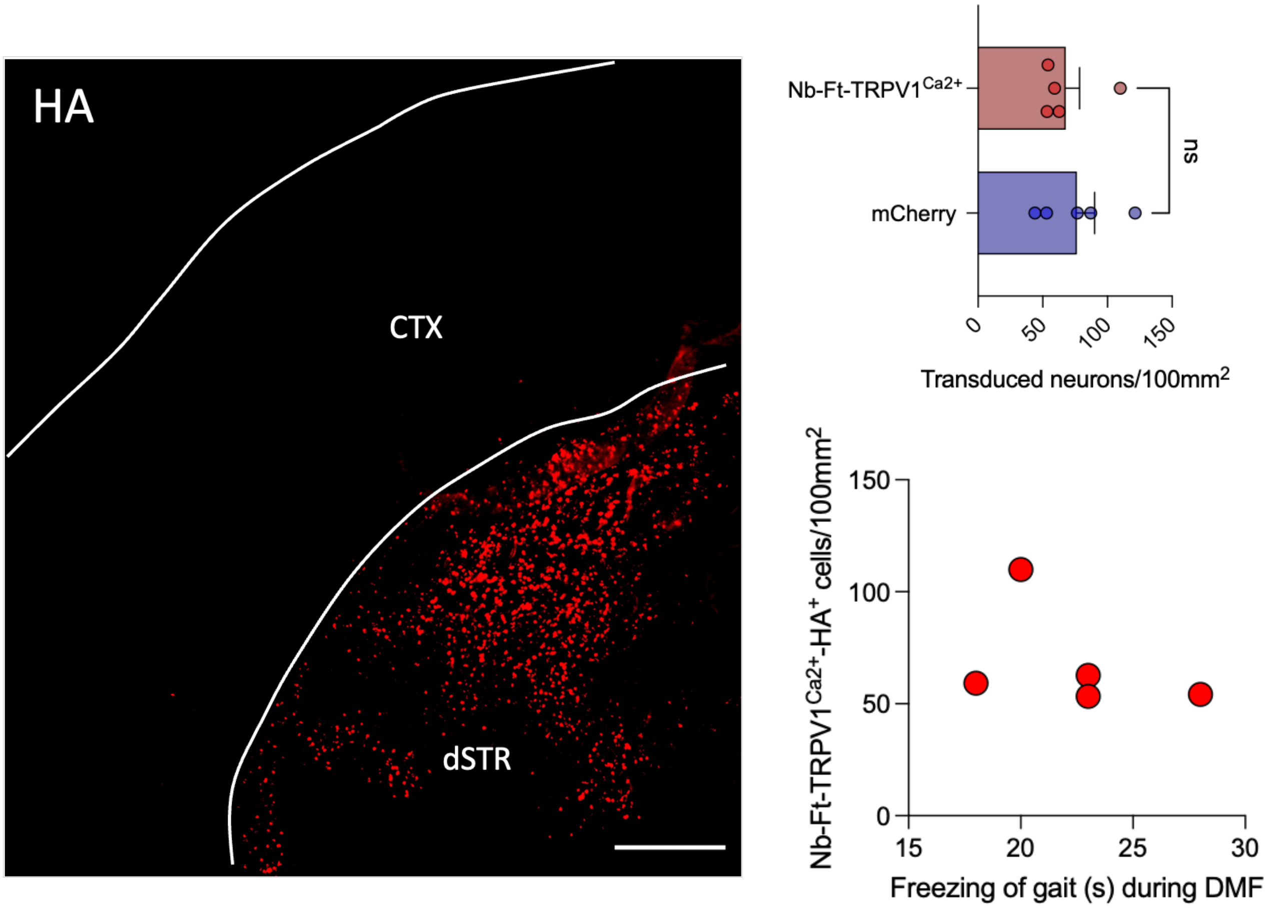

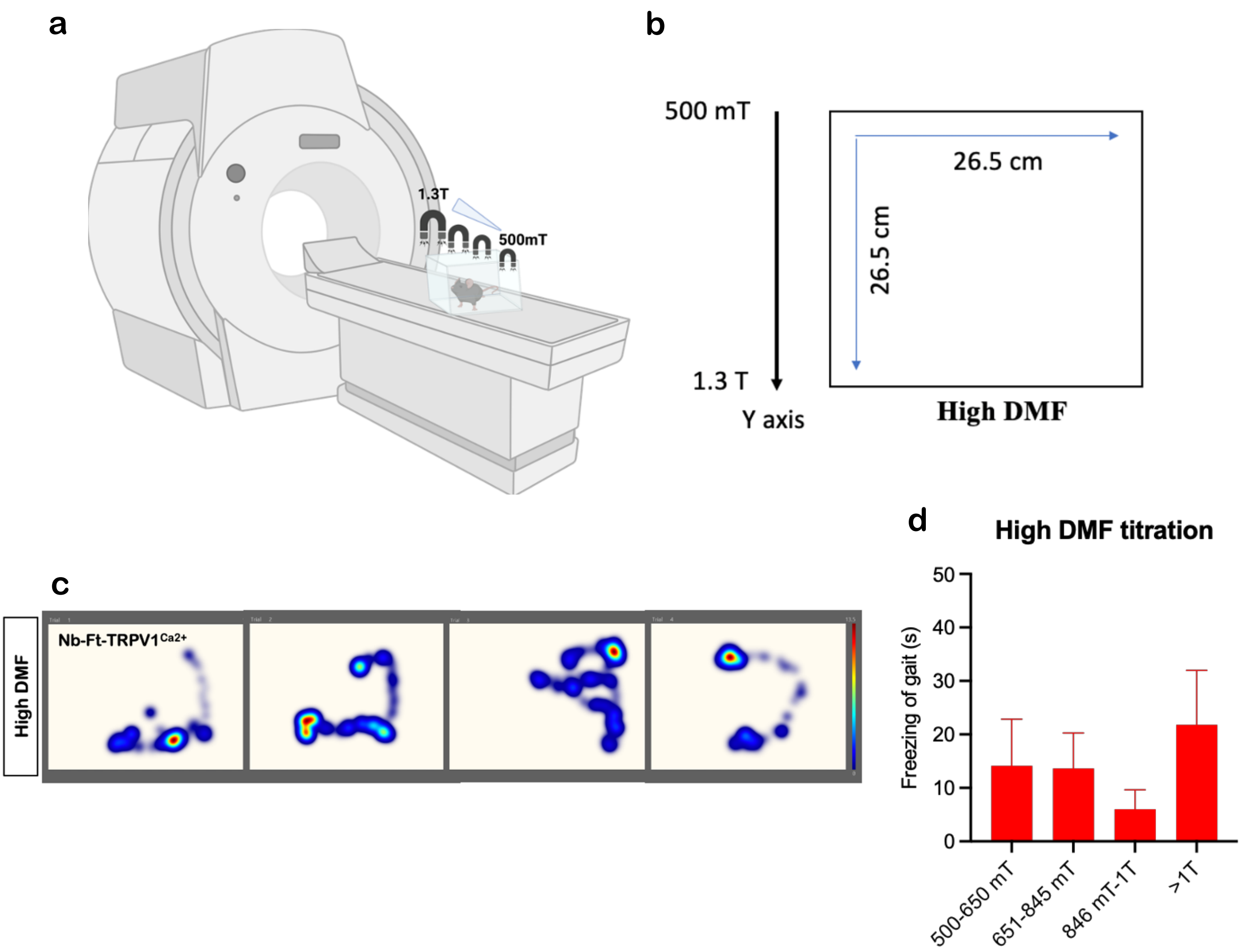

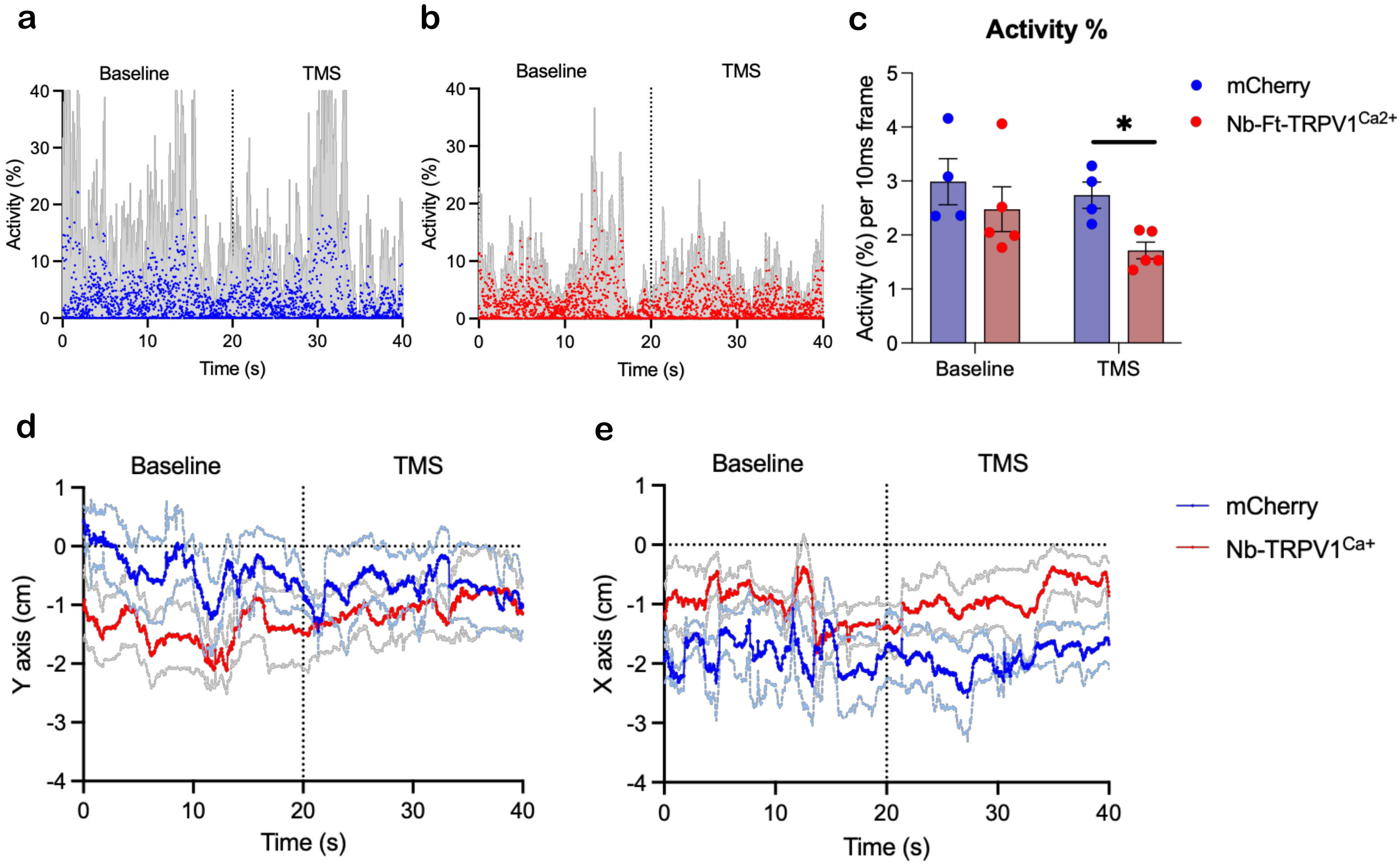

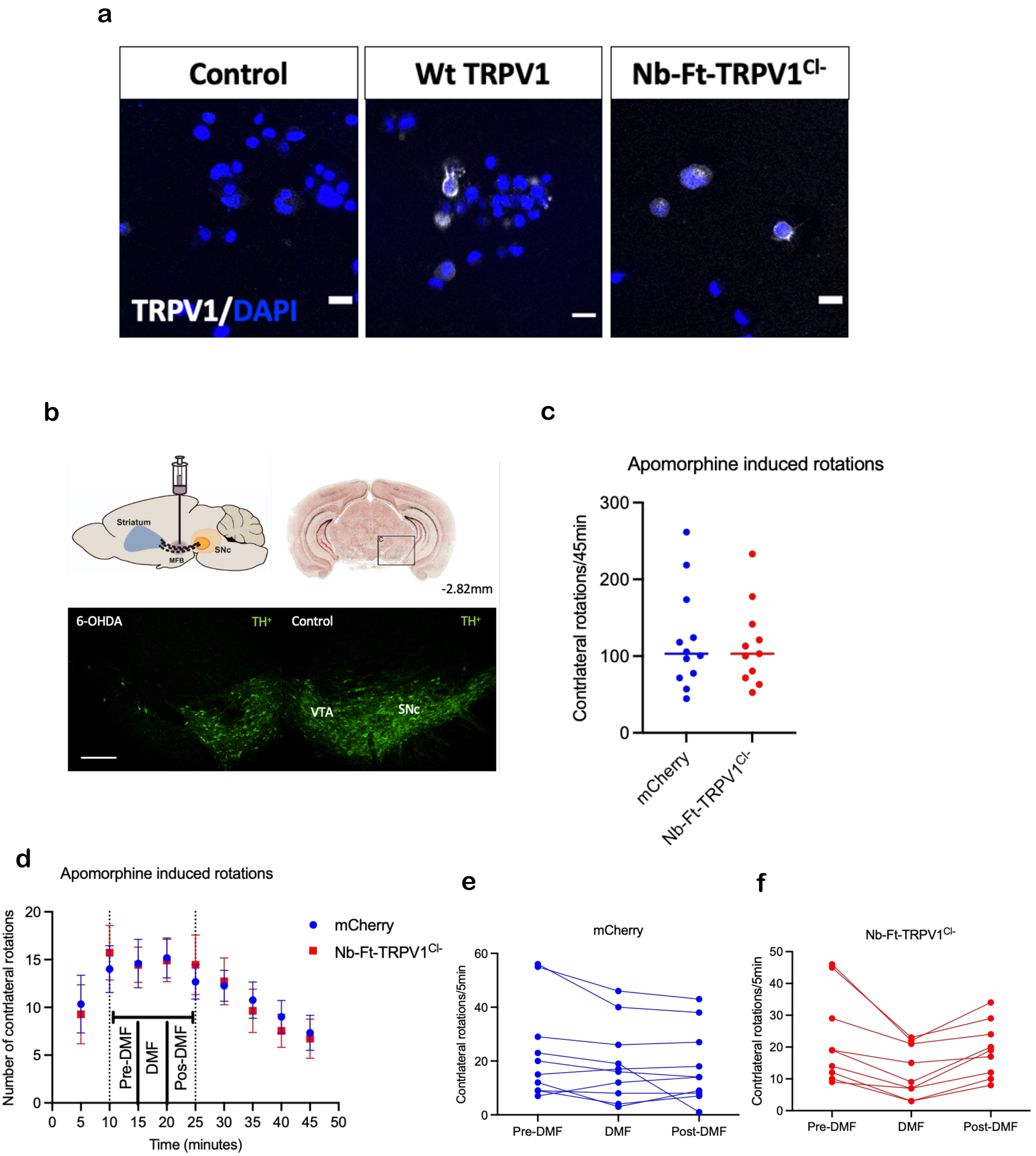
